# Diffusion of small molecule drugs is affected by surface interactions and crowder proteins

**DOI:** 10.1101/2021.12.30.474528

**Authors:** Debabrata Dey, Ariane Nunes-Alves, Rebecca C Wade, Gideon Schreiber

**Affiliations:** Department of Biomolecular Sciences, Weizmann Institute of Science, Israel; Molecular and Cellular Modeling Group, Heidelberg Institute for Theoretical Studies, Schloss-Wolfsbrunnenweg 35, 69118 Heidelberg, Germany; Center for Molecular Biology (ZMBH), DKFZ-ZMBH Alliance, Heidelberg University, Im Neuenheimer Feld 282, 69120 Heidelberg, Germany; Interdisciplinary Center for Scientific Computing (IWR), Heidelberg University, Im Neuenheimer Feld 205, Heidelberg, Germany.; Institute of Chemistry, Technische Universität Berlin, 10623 Berlin, Germany

## Abstract

Crowded environments are known to affect the diffusion of macromolecules but their effects on the diffusion of small molecules, such as drugs, are largely uncharacterized. Here, we investigate how three macromolecular protein crowders, bovine serum albumin (BSA), hen egg-white lysozyme and myoglobin, influence the translational diffusion rates and interactions of four low molecular-weight compounds: the diagnostic marker fluorescein, and three drugs, doxorubicin, glycogen synthase kinase-3 inhibitor SB216763 and quinacrine. Using Fluorescence Recovery After Photo-bleaching in Line mode (Line-FRAP), Brownian dynamics simulations and molecular docking, we find that the diffusive behavior of the small molecules is highly affected by self-aggregation, interactions with the proteins, and surface adhesion. The diffusion of fluorescein is decreased by protein crowders due to its interactions with the proteins and their surface adsorption. In contrast, the presence of protein crowders increases the diffusion rate of doxorubicin by reducing surface interactions. SB216763 shows a third scenario, where BSA, but not lysozyme or myoglobin , reduces self-aggregation, resulting in faster diffusion. Quinacrine was the only compound whose diffusion was not affected by the presence of protein crowders. The mechanistic insights gained here into the effects of interactions of small molecules with proteins and surfaces on the translational diffusion of small molecules can assist in optimizing the design of compounds for higher mobility and lower occlusion in complex macromolecular environments.

## Introduction

Most drugs have a molecular weight < 1 kDa and an octanol/water partition (logP) of 1– 6 (1). For a drug to be orally absorbed, it must be hydrophobic to partition into a lipid bilayer, but not to such an extent that it will result in permanent absorbance into the bilayer (1). This property results in drugs being able to self-associate as well as bind to hydrophobic cellular components through soft interactions (2), which in turn decreases their active concentration and affects their diffusion. In addition to self-aggregation in polar aqueous or buffer-like solvents, small molecule drugs tend to interact with or adsorb to glass or plastic surfaces, further complicating diffusion measurements (3). Overlooking these factors may lead to spurious observations that can negatively impact studies of small molecule drugs (4, 5). Moreover, while biophysical characterization of such drugs is carried out *in vitro*, they are required to be active in a complex, crowded environment, containing membranes and macromolecules at concentrations of up to 300 mg/mL (6).

For small molecules to reach their target within a crowded milieu, they have to freely diffuse and avoid off-target interactions (7). Passive diffusion is considered to be a primary mechanism of intracellular drug transport (8). However, more recently carrier-mediated transport (8, 9) or a combination of the two were also identified (10). The study of the diffusion of small organic molecules inside complex environments is therefore highly relevant for drug design (7, 11), as well as for soft matter and biochemical studies more generally (12) (13). The challenge (14) is that most small molecules are not fluorescent in the visible spectrum, making it almost impossible to follow them by optical methods (15). Conversely, proteins can usually be fluorescently labeled without significant perturbation of their diffusion rate and function, making it relatively easy to study their diffusion (16).

Here, we investigated the diffusion of three fluorescent organic small molecule therapeutic drugs and one diagnostic marker in solutions containing proteins as crowders. Doxorubicin (DOX) is widely used as an anti-cancer agent (17); quinacrine dihydrochloride has been used for antimalarial and antiprotozoal therapy (18); glycogen synthase kinase-3 (GSK3) inhibitor SB216763 is a potent selective and ATP-competitive inhibitor of glycogen synthase kinase-3 and significantly prevents lung inflammation and fibrosis in mouse models (19). DOX and quinacrine behave as weak bases at physiological pH (20). Unlike these three therapeutic drugs, fluorescein in its salt form, which is widely used as a biological marker in angiographic assays, is a negatively charged molecule at physiological pH (21). These four molecules were chosen for study as they form a structurally and chemically diverse set of compounds that fluoresce in the visible spectrum, a prerequisite for Line FRAP studies (Figure 1). The diffusion and aggregation of these low MW molecules were evaluated in buffer, and with bovine serum albumin (BSA), hen egg-white lysozyme (HEWL), or myoglobin as molecular crowders (22, 23). Self-aggregation and binding to these proteins can potentially mimic their behaviors in body fluids or inside the cell (24). Diffusion rate measurements provide a good means of estimating these protein-ligand associations or the aggregation state of the compounds in diverse environments which affect the mobility of the compounds.

**Figure 1:**
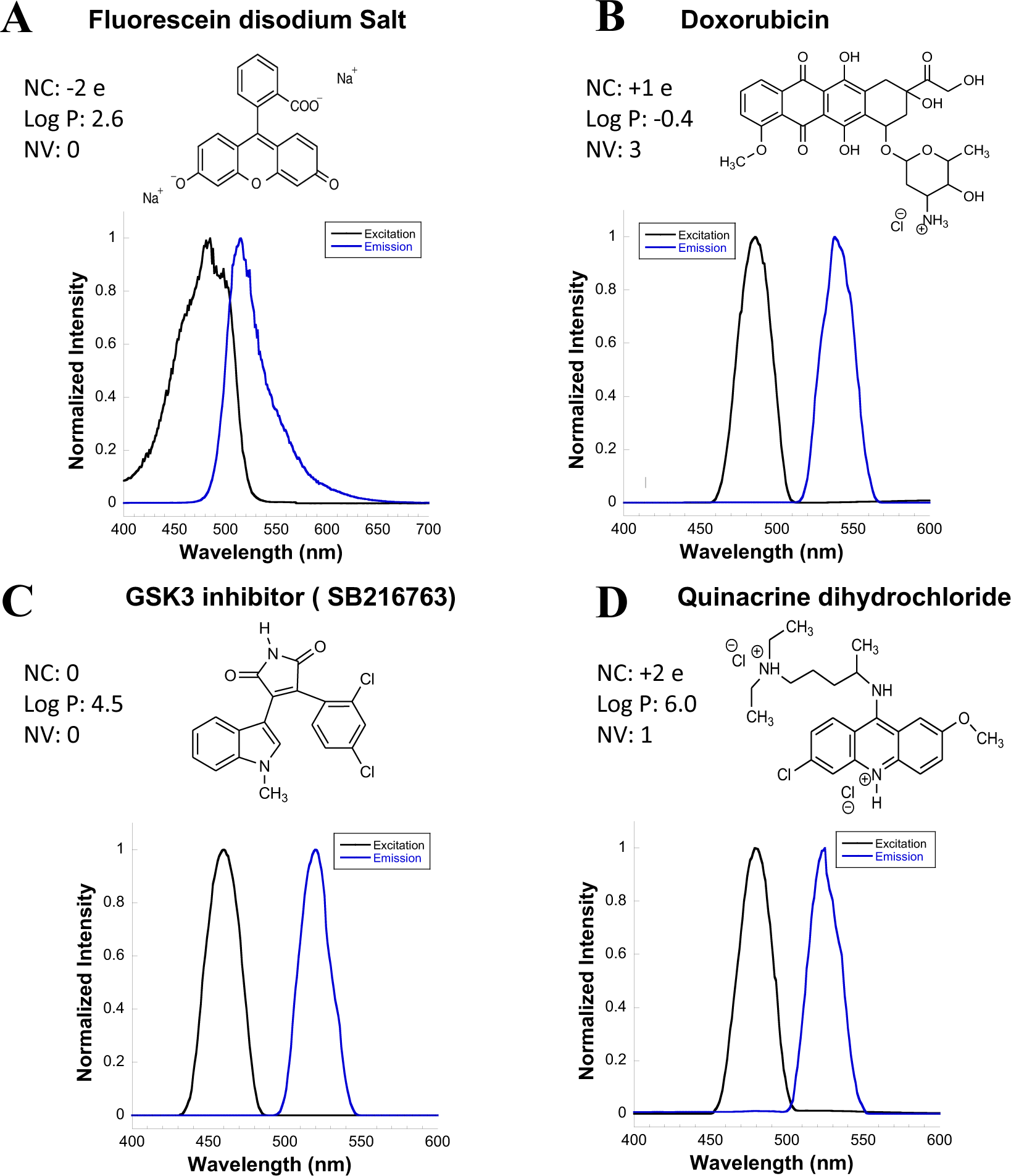
Chemical structures, physicochemical properties and comparative excitation and emission spectra with relative strengths: (A) fluorescein disodium salt, (B) doxorubicin, (C) GSK3 SB216763 inhibitor, and (D) quinacrine dihydrochloride. NC: net charge at pH 7.2 (computed using Epik (51)). Log P: log of the octanol-water partition coefficient (computed using QikProp (52)). NV: number of violations to Lipinski’s Rule of Five for drug likeness (computed using QikProp (52)).

A wide range of computational and experimental techniques have been developed to compute and measure diffusion rates, as well as to study molecular association. On the computational side, Brownian dynamics (BD) (25) and molecular dynamics (MD) (14, 23, 26, 27) simulations are widely used. In both methods, the system can be considered in atomic detail, but in BD simulations, the molecules are generally treated as rigid bodies and an implicit solvation model is used. These simplifications allow BD simulations to routinely achieve tens of microseconds of simulation time with systems containing hundreds of solute molecules. BD simulations have been previously employed to compute diffusional properties of proteins, revealing, for example, molecular details about the reduction in diffusion rates of BSA, myoglobin, hemoglobin and γ-globulin under crowding conditions (28, 29) or the adsorption of HEWL to an inorganic surface (30, 31) . Among the experimental techniques, fluorescence recovery after photobleaching (FRAP) (32), fluorescence correlation microscopy (FCS) (33), and single-particle tracking (SPT) (34) are the most popular. Each method has its benefits and limitations (35). FCS is considered the gold standard for this purpose; however, its application can be challenging. Firstly, it requires high quantum yield (which is rare for drugs), and secondly, it can be performed only at very low concentrations, which may be much lower than found *in vivo*. Therefore, FRAP is the technique most widely used by experimental biologists (35). It is fast, non-invasive, highly specific, and relatively easy to perform (35). Moreover, FRAP can be used for molecules with poor quantum yields and at high, biologically relevant concentrations (32).

We previously developed the Line-FRAP method to monitor the diffusion rates of proteins in various environments (36). The main advantage of Line-FRAP over conventional FRAP is the much faster data acquisition rate, which allows measurements for fast diffusing molecules. The apparent diffusion coefficients derived from FRAP measurements are D_confocal_ (36), which are calculated according to equation 1:

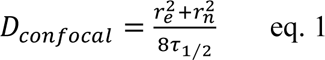

where 1_1/2_ is the half-time of recovery and r_e_ and r_n_ are the effective and nominal bleach radii. Line-FRAP D_confocal_ values were shown to be in line with FCS diffusion rates for proteins (36). However, for small molecules, they provide only relative 3D diffusion coefficients (37). An alternative way to present FRAP results is by reporting only τ_1/2_ values. However, as seen from eq. 1, this completely ignores the contribution from the bleach radius (which is squared) (38, 39). In particular, r_e_ varies, depending on the type of molecule and the diffusion condition, making its measurement critical to obtain reliable estimates of diffusion rates.

In this study, we determined D_confocal_ values by Line-FRAP for the four low molecular weight compounds using the three different proteins as crowders. The derived diffusion coefficients are found to depend on the self-aggregation properties of the small molecules, their binding to proteins and surfaces, and the buffer solution conditions. The protein crowders affect the solubility and diffusion rates of the compounds in a manner that is dependent on the properties of the different protein crowders used. Complementary steady-state fluorescence quenching and size exclusion chromatography experiments provide thermodynamic data that reveal differences in the association between the compounds and the protein crowders. Furthermore, Brownian dynamics simulations and molecular docking for the systems studied experimentally shed light on the intermolecular interactions and molecular mechanisms responsible for the differences in the diffusion coefficients measured for the different small molecule and protein crowder combinations.

## Results

We used high-content screening of over 1000 drugs and the database fluorophores.org to search for drugs that are fluorescent in visible light, allowing us to follow their diffusion using FRAP. Of those identified, we chose fluorescein, DOX, SB216763, and quinacrine for this study, as they have moderate to good quantum yields.

### Fluorescein disodium salt

Fluorescein is a negatively charged organic small molecule (MW = 376.3 Da, Figure 1A). We first measured the diffusion rates of fluorescein in different solvents with and without protein crowders. The values of D_confocal_ in DMSO and PBS, with and without Tween20, are given in Figure 2A and Figure 2B shows micrograph images of fluorescein in corresponding solutions. A D_confocal_ value of ∼56 µm^2^s^-1^ was determined in all these cases, and there was no evidence for aggregation of fluorescein. Next, we measured fluorescein’s diffusion in the presence of increasing concentrations of BSA, HEWL, and myoglobin (Figure 2 C-E). We were surprised to see that, even at low concentrations of protein (5-20 mg/mL), the presence of BSA, HEWL and myoglobin slowed down the diffusion of fluorescein significantly (Figure 2E). Since the fraction of excluded volume due to the presence of protein crowders at these concentrations is low, we suspected that quinary or weak interactions between fluorescein and the protein crowders are the main drivers of the reduction in the diffusion rates of fluorescein, in line with previous publications (40). HEWL had the biggest effect on diffusion, followed by BSA and myoglobin. Interestingly, for myoglobin, increasing protein concentrations above 5 mg/mL resulted in an increase in diffusion rates, which returned to their level without crowder at 50 mg/mL (Figure 2E). As in the absence of protein crowders, micrographs in the presence of protein crowders did not show aggregation (Figure 2D) and FRAP fully recovered after bleach under all conditions measured (Figure 2C).

**Figure 2:**
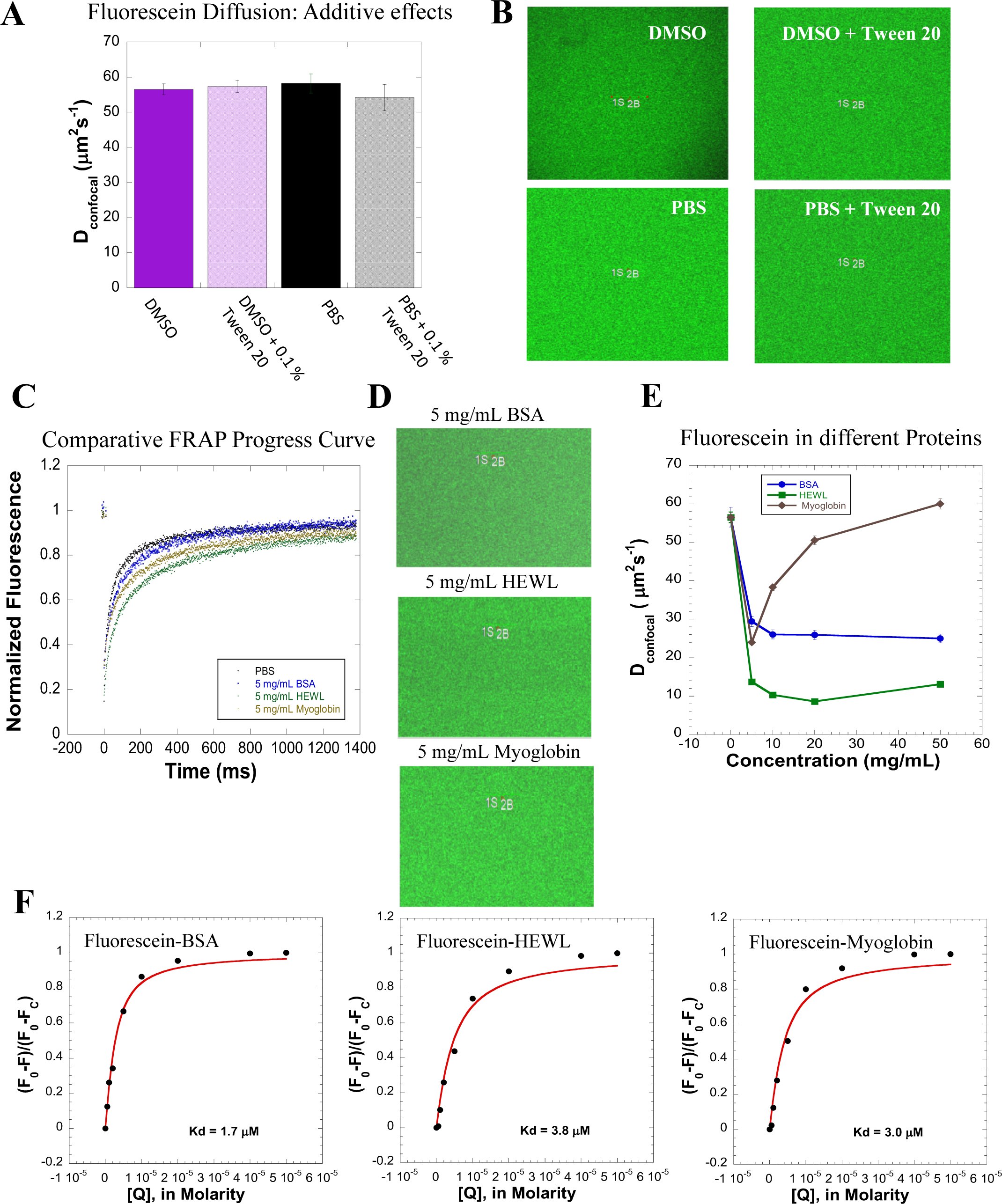
Fluorescein in buffer and the presence of protein crowders: **(A,B)** Diffusion coefficients (A) and images from confocal microscopy (B) of fluorescein in DMSO or PBS with or without Tween 20. **(C,D)** Comparative averaged FRAP profiles ( N= 30; R= 0.99 for each of the fits) (C) and images from confocal microscopy (D) of fluorescein in the presence of 5 mg/mL BSA, HEWL or myoglobin. **(E)** Diffusion coefficients of fluorescein from measurements at increasing concentrations of the three protein crowders. **(F)** Determination of binding affinities of fluorescein to the three protein crowders by fluorescence quenching in PBS buffer. (F_0_-F)/(F_0_-F_c_) vs [Q] plots of the data and fits (R=0.99), where [Q] is the titrating drug concentration in molarity, are shown. Micromolar binding affinities were determined for fluorescein to all three protein crowders.

Next, we compared the diffusion of fluorescein and labeled BSA (Figure 3D) to test if the reduction in the diffusion rates of fluorescein in the presence of BSA could be explained solely by interactions with BSA in solution. Unexpectedly, we observed that the diffusion coefficients of labeled BSA were higher than those of fluorescein in the presence of BSA for all concentrations of BSA tested. This observation implies that the reduced diffusion coefficient of fluorescein in the presence of BSA cannot be solely explained by binding of fluorescein to freely diffusing BSA.

**Figure 3:**
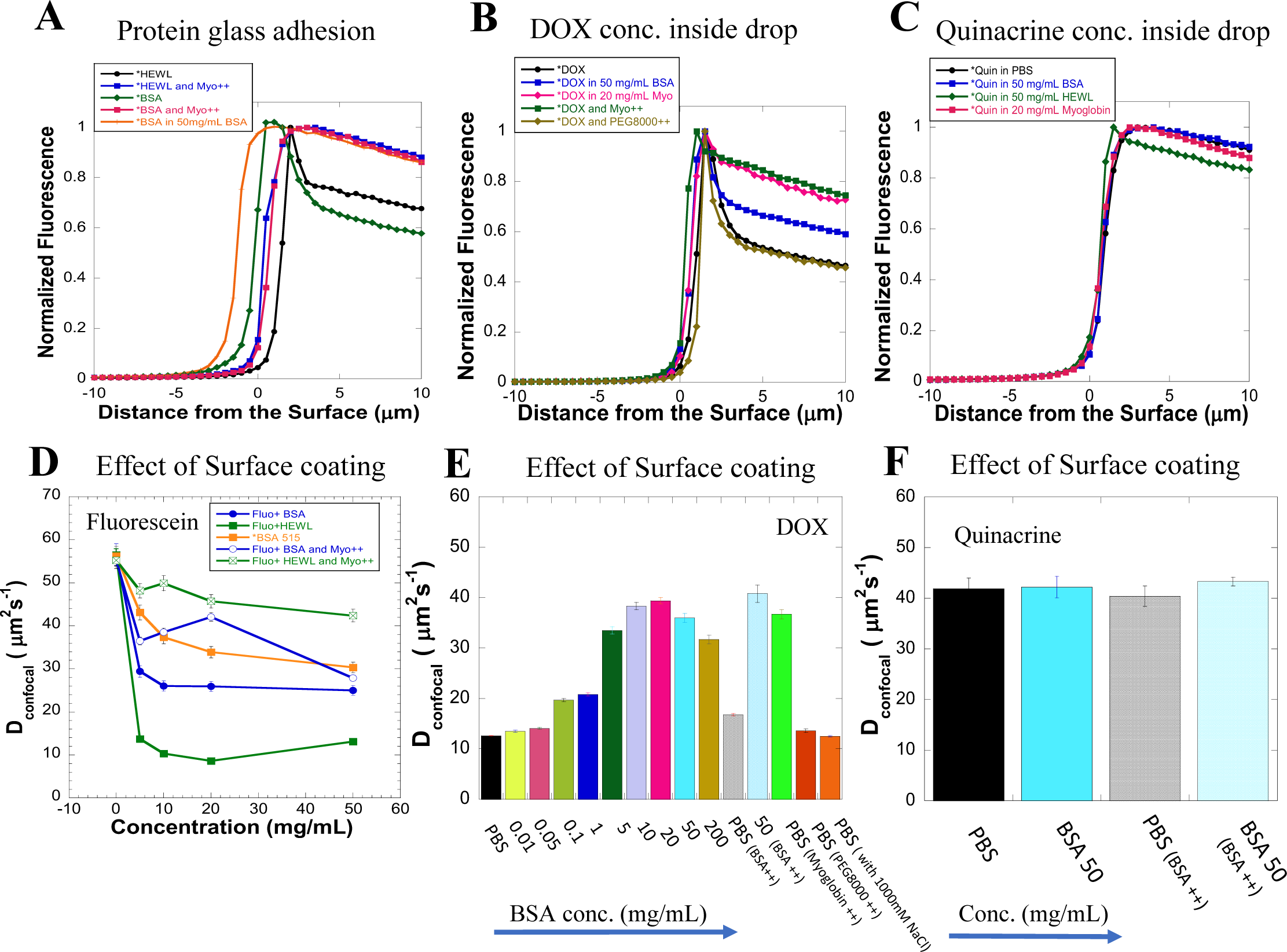
Adsorption of drugs and protein crowders to a glass surface. **(A-C)** Determination of drop homogeneity of (A) labelled proteins (protein-glass adhesion), (B) doxorubicin, and (C) quinacrine in different crowding environments and in buffer solutions with or without surface coating. **(D-F)** Effect of surface coating of glass plates by BSA or myoglobin on the diffusion coefficients of the small molecules for (D) fluorescein, (E) doxorubicin and (F) quinacrine. Here starting with * symbol in the figure legends denotes fluorophore either labelled with proteins or small molecule which are itself fluorescent and ending with ++ symbol denotes the glass coating.

However, these experiments were performed on liquid drops on a glass surface and, if BSA was adsorbed to the glass surface, this could lead to a reduction in the diffusion of BSA-bound fluorescein molecules. To directly assess this possibility, a drop containing labeled BSA or labeled HEWL was applied to the glass, and the fluorescence along the z-axis perpendicular to the plane of the glass surface was measured (Figure 3A). Clearly, both proteins attach to the surface, as seen by the higher fluorescence close to the surface. Next, we pre-coated the glass slides either with unlabeled BSA or with myoglobin, washed the glass, and then applied labeled BSA or HEWL (Figure 3A). Now, the fluorescence profile indicated that the labeled protein became rather homogeneously distributed along the z-axis above the surface, indicating lack of adsorption of labeled protein to the glass. Repeating the FRAP measurements of fluorescein in the presence of either HEWL or BSA, but this time after pre-coating the glass with myoglobin (Figure 3D), resulted in much higher diffusion coefficients for fluorescein. In the presence of increasing concentrations of BSA, the D_confocal_ values for fluorescein were similar to those measured for labeled BSA. Furthermore, after coating the surface with myoglobin, the presence of HEWL had only a small effect on the diffusion coefficient of fluorescein, in contrast to the large reduction in D_confocal_ without pre-coating. As fluorescein alone in PBS or in the presence of protein crowders does not attach to the glass surface (Figures 3A and S7A), the experimental data suggest that fluorescein’s reduced diffusion is due to its attachment to the proteins bound to the glass surface. This conclusion is supported by dynamic light scattering (DLS) experiments to measure the hydrodynamic size of BSA alone and in the presence of fluorescein at different protein and fluorescein concentrations (Figure S8). The hydrodynamic size (in nm) of BSA in did not change on addition of fluorescein to different concentrations of BSA or even when the added fluorescein concentration was 10 times that used in the FRAP measurements. This shows that fluorescein does not affect the oligomerization state of BSA and that its interaction with freely diffusing BSA would be expected to give a diffusion coefficient corresponding to that of BSA, which is higher than observed in the experiments.

To quantify the protein-small molecule binding affinities, steady-state fluorescence quenching experiments were carried out. The association of the small molecules with the proteins causes a change in the environment around buried tryptophan residues (which are largely responsible for the intrinsic fluorescence properties of proteins), which results in the quenching of fluorescent signals from the protein (41). For example, BSA contains two tryptophan residues, Trp-134 and Trp-212, located in the first and second domains of hydrophobic protein regions (41). The decrease of fluorescence intensity for BSA was monitored at 344 nm wavelength for the drug-protein pairs. Figures S6 A-C show representative fluorescence quench spectra for fluorescein with BSA, HEWL, and myoglobin. We assume that the observed changes in the fluorescence are the result of the interaction between the small molecule and the protein. Therefore, corresponding plots of (F_0_-F)/(F_0_-F_c_) versus fluorescein concentration [Q] in molarity (Figure 2F) were used to determine the affinity between them, using equation 2 (see Materials and methods). BSA, HEWL, and myoglobin were found to bind fluorescein with affinities of 1.7±0.2 µM, 3.8±0.6 µM and 3.0±0.5 µM, respectively (Figure 2F and Table S1).

To further investigate the factors influencing the diffusion of fluorescein, BD simulations were performed for fluorescein and the same concentrations of protein crowders as present in FRAP experiments. The results show that the effects of the crowders modelled in these simulations (excluded volume, electrostatic and hydrophobic interactions between rigid solutes) result in modest reductions of up to 15% in the computed translational diffusion coefficients of the small molecules at crowder concentrations up to 50 mg/mL (Figure S1). This is roughly in line with the reduction in D_confocal_ observed for fluorescein and HEWL and BSA crowders after pre-coating the glass slide with myoglobin. The BD simulations also revealed differences in the interactions of fluorescein with the three protein crowders which were examined by computing the number of intermolecular contact interactions. Contacts were defined as present if non-hydrogen atoms (at least one in a protein crowder and at least one in a small molecule) were within 4.5 Å of each other. This distance was chosen to capture electrostatic and van der Waals interactions between protein crowders and fluorescein. The number of protein- fluorescein contact interactions (Figures 4A) and the peak of the radial distribution function (RDF) for protein-small molecule distances (Figures 4B) are higher in the presence of HEWL, showing that the interactions of fluorescein are stronger with this protein crowder. The stronger interactions between fluorescein and HEWL are consistent with the experimentally observed stronger reduction in the diffusion coefficient of fluorescein in the presence of HEWL than in the presence of the other two protein crowders, without pre-coating the glass slides (when fluorescein can interact with the slower diffusing HEWL on the glass surface). The stronger interaction between fluorescein and HEWL can be explained by the strong electrostatic interaction between them, since at the pH of the experiments (7.4), fluorescein is negatively charged (-2e) and HEWL is positively charged (+8e). Another factor that contributes to this strong electrostatic interaction is the distribution of the molecular electrostatic potential of HEWL, which has a large positively charged region on the surface that interacts mostly with fluorescein during the BD simulations (Figure S5). Despite their negative net charges, myoglobin (-2e) and BSA (- 16e) can also make favorable electrostatic interactions with fluorescein through regions of positive electrostatic potential on their surfaces and these are the parts of the protein surface with the greatest occupation of contacts with fluorescein during the BD simulations (Figure S5). However, these interactions are weaker than for HEWL and result in fewer contacts and much less pronounced peaks in the RDF (Figures 4, S3 and S4).

**Figure 4:**
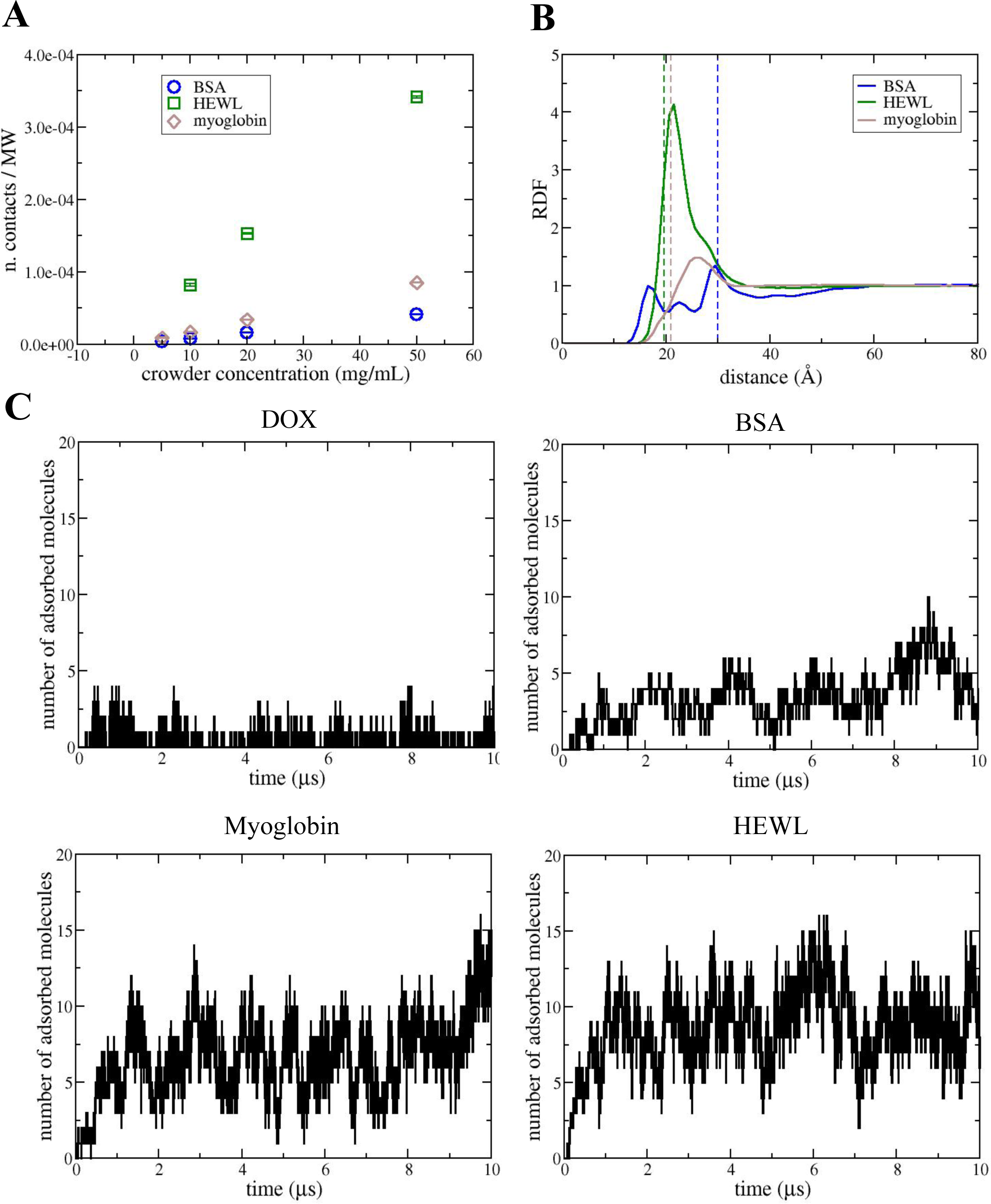
BD simulations reveal protein-drug interactions, and protein or drug adsorption to a silica surface. Interactions between fluorescein and protein crowders in BD simulations show differences between the three protein crowders. The number of protein-fluorescein contact interactions and the RDF peak are higher in the presence of HEWL, indicating stronger interactions of fluorescein with this protein crowder. (A) Number of protein-fluorescein contact interactions divided by the molecular weight (MW) of the protein crowder (66637, 17820 and 14331 g/mol for BSA, myoglobin and HEWL, respectively) plotted against protein crowder concentration. (B) Radial distribution functions (RDF) for protein-fluorescein distances computed from simulations with 50 mg/mL of crowder (pH 7.2, ionic strength of 190 mM). The dashed lines indicate the sum of the Stokes radii of the protein crowder and of fluorescein. Stokes radii of proteins: 25.7 Å, 16.6 Å and 15.3 Å for BSA, myoglobin and HEWL, respectively. Stokes radius of fluorescein: 4.3 Å. (C) Number of doxorubicin or protein crowder molecules adsorbed to a silica surface in BD simulations performed with one type of solute molecule (80 molecules of DOX or 440 molecules of protein crowder at 50 mg/mL concentration, pH 7.2, ionic strength of 190 mM). The numbers were obtained from a single BD simulation performed for 10 microseconds. The region adjacent to the surface was initially depleted of solute molecules (see Methods for details). The silica surface is negatively charged and thus electrostatic interactions favor the binding of HEWL (+8 e) relative to BSA (-16 e).

In summary, both experiments and simulations show that fluorescein makes contacts with all three crowders, with the highest number of contacts observed for binding to HEWL. This results in a modest reduction of the diffusion rates. However, the three proteins are adsorbed to the glass surface, with HEWL being the strongest adsorbed. This results in further reduction in fluorescein diffusion rates, as fluorescein interacts both with the immobile (surface adsorbed) and the mobile (in solution) protein molecules, with the reduction being greatest in the presence of HEWL.

### Doxorubicin

The chemical structure and spectroscopic properties of DOX (MW=580 Da) are shown in Figure 1B. It diffuses surprisingly slowly in PBS buffer, with D_confocal_∼ 12.6 µm^2^s^-1^ (Figure 5A), while the infinite dilution translational diffusion coefficient of DOX (Stokes radius of Å), D_trans_, is calculated to be 412 µm^2^s^-1^ (Table S2). Next, we measured the diffusion of DOX in DMSO, where D_confocal_ increased to 47 µm^2^s^-1^ (Figure 5A). Addition of 0.1 percent Tween 20 surfactant, which promotes the solubilization of aggregated hydrophobic DOX molecules and prevents surface attachment (3), increased D_confocal_ in PBS to 27 µm^2^s^-^1, with a similar value measured with 10 mg/mL BSA, with or without Tween 20 (Figure 5A). Micrographs of DOX in different solutions (Figure 5A, right panel) do not show obvious high MW aggregates. However, this does not exclude that DOX forms lower MW aggregates, as has been suggested previously (42), and could explain the slow diffusion of DOX in PBS. To further verify the D_confocal_ values, we measured them after 10 and 63 ms bleach times. The corresponding recovery curves and bleach sizes are shown in Figure S9. The data clearly show much slower recovery curves after 63 ms bleach, which are accompanied by a wider bleach radius. However, the D_confocal_ values calculated from τ_1/2_ and r_e_ were similar for the different bleach times (Table S3), in line with what we have previously found for protein diffusion (36).

**Figure 5:**
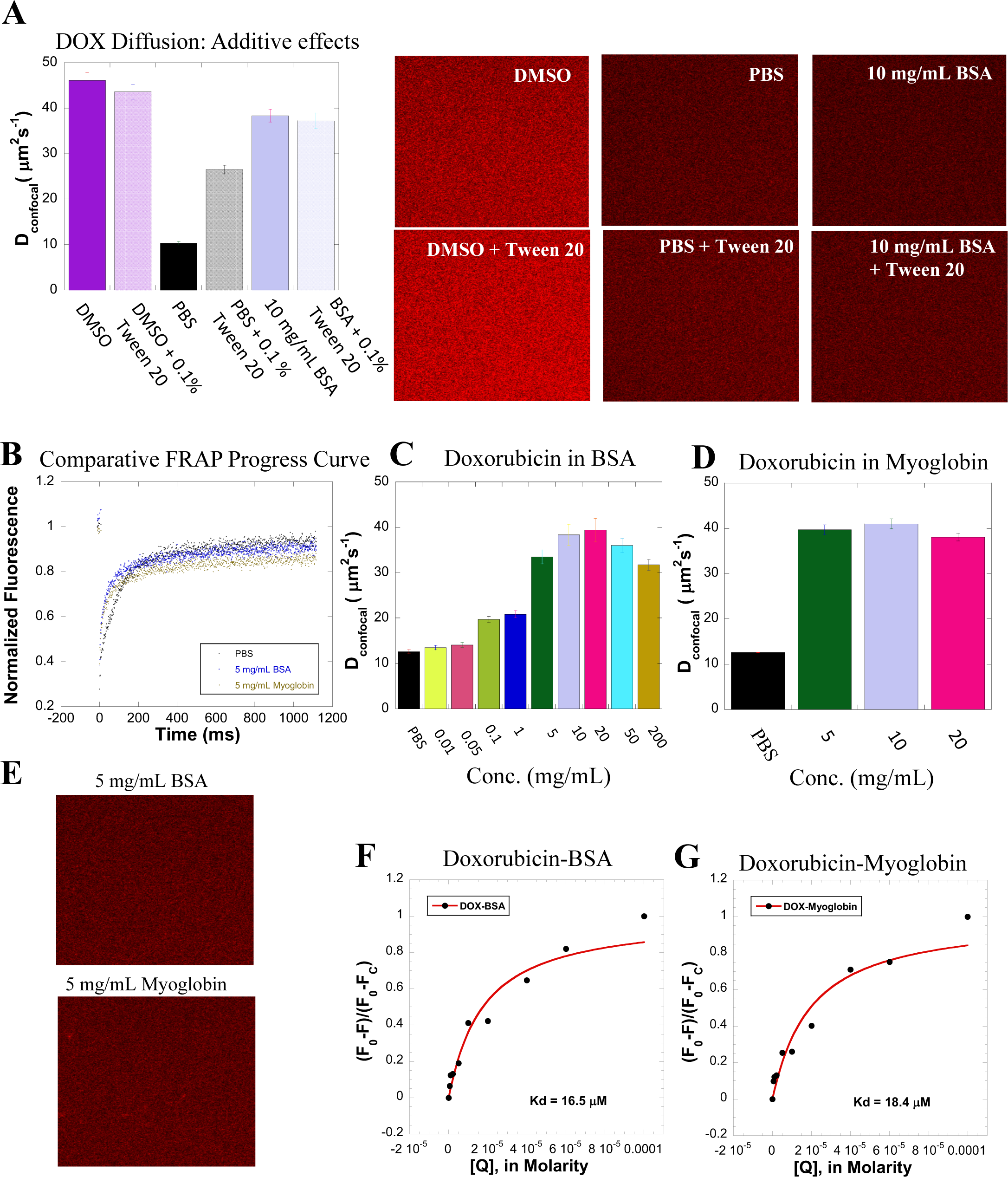
Doxorubicin diffusion in protein crowders. (A) Diffusion coefficients and images from confocal microscope of doxorubicin in DMSO, PBS or BSA solutions with or without Tween 20. (B) Averaged FRAP profiles (N= 30; R= 0.99 for each of the fits) in PBS and in the presence of 5 mg/mL BSA and 5 mg/mL myoglobin. (C, D) Diffusion coefficients of DOX in increasing amounts of (C) BSA and (D) myoglobin. (E) Confocal microscopy images of doxorubicin in the presence of 5 mg/mL BSA and 5 mg/mL myoglobin . (F, G) The results of binding affinity measurements showing (F_0_-F)/(F_0_-F_c_) vs [Q] plots from fluorescence quenching experiments of (F) DOX-BSA (R= 0.98), (G) DOX-myoglobin (R= 0.97) systems in PBS, where Q is the titrating drug concentration in molarity.

Next, we measured DOX diffusion after addition of different concentrations of BSA or myoglobin (measurements with HEWL crowders were not possible as DOX precipitated on addition of HEWL). The addition of these proteins increased D_confocal_ values up to ∼ 40 µm^2^s^-1^ (Figures 5 C, D), close to the value recorded in DMSO (Figure 5A). Titration of increasing concentrations of BSA, from 0.01-200 mg/mL, showed that the maximum diffusion coefficient is reached at 10-20 mg/mL, with a decrease observed at higher BSA concentrations. This reduction can be associated with the excluded volume effect, which starts to be significant for BSA at the higher concentrations measured (fraction of occupied volume of 0.035 at 50 mg/mL BSA and 0.14 at 200 mg/mL BSA). BD simulations (which include intermolecular forces as well as excluded volume effects) show that not only the diffusion coefficient of DOX, but also that of BSA itself, is reduced as BSA concentrations increase from 20 to 200 mg/mL (Figures S1 and S2), in agreement with the experimental results.

Next, we determined the binding of DOX to BSA and myoglobin by fluorescence quenching titration experiments (Figures 5F and G & S6 D and E). BSA and myoglobin bind DOX with low affinities of 16.5±2.5 µM and 18.4±3.1 µM, respectively (Figure 5F and G and Table S1), in agreement with a previous study (41).

To investigate whether the slow diffusion of DOX in PBS is a result of surface attachment or self-aggregation, we determined the fluorescence intensity as a function of the distance from a glass surface for DOX. Whereas for fluorescein, we saw a homogeneous distribution of the small molecule in the drop, for DOX, surface attachment was clearly observed (Figures S7A-B and 3B). This is in line with a previous report that DOX can adsorb to a polypropylene surface (3). We then used BSA, myoglobin and PEG8000 solutions to coat the glass surface and thereby decrease the surface attachment and adsorption of the DOX molecules. Out of the three, myoglobin acted as the best surface coating agent for DOX molecules (Figure 3B), resulting in almost complete removal of DOX from the surface. In addition, myoglobin coating of the glass surface resulted in a large increase in D_confocal_ compared to DOX in PBS solution (Figure 3E), giving a value of 36 µm^2^s^-1^, which is similar to the maximum value measured in the presence of BSA. The effect of BSA or PEG8000 coating of the glass surface on the D_confocal_ values was minimal and DOX remained attached to the glass surface even after coating (Figure 3B and E). BD simulations in the absence of a surface show a low number of DOX-protein contacts (Figure S3) and small, broad RDF peaks for DOX-protein distances (Figure S4). This agrees with the experimental results, which show that the main effect of the protein crowders, especially at low concentrations, is prevention of DOX attachment to the glass surface, rather than slowed diffusion by quinary interactions with protein crowders, as observed for fluorescein. Nonetheless, it should be noted that quinary interactions between DOX and BSA are expected, as reported in a previous study (41).

BD simulations performed with a silica surface to mimic the glass plate and DOX molecules or the protein crowders at a concentration of 50 mg/mL show that all these types of molecules can adsorb to the negatively charged silica surface (Figures 4C), in agreement with previous studies using simulations or experiments (30, 43–45). However, a higher number of myoglobin and HEWL molecules adsorbed to the surface compared to DOX and BSA, indicating that myoglobin and HEWL interact better with the surface and may be better at preventing DOX surface attachment, consistent with the experimental results observed for DOX with myoglobin and BSA crowders. Measuring diffusion of DOX in PBS solution with an increased ionic strength (1000 mM NaCl) resulted in D_confocal_ of 12.5 µm^2^s^-1^ (Figure 3E), suggesting that the electrostatic attraction of the positively charged DOX to the negatively charged glass surface is not required for surface adsorption which is instead driven by hydrophobic interactions, in line with a previous report of DOX adsorption to polypropylene surfaces (3).

### GSK3 inhibitor SB216763

GSK3 is a protein kinase that is active in a number of central intracellular signaling pathways, including cellular proliferation, migration, glucose regulation, and apoptosis (19). The GSK3 inhibitor SB216763 is currently being evaluated for several related malignancies (19). The chemical structure and spectroscopic properties of SB216763 are shown in Figure 1C. SB216763 has a strong tendency to aggregate in PBS (Figure 6A). Aggregation is also observed in DMEM media. Of the four small molecules tested, the GSK3 inhibitor is the only one with neutral net charge at the pH of the experiments. The lack of repulsive electrostatic interactions and the high hydrophobicity (indicated by a logP value of 4.5, Figure 1C) may facilitate aggregation. Moreover, BD simulations of the GSK3 inhibitor without protein crowders showed greater self-interactions than those observed for the other small molecules (Table S4), consistent with the experimental data. The addition of BSA (but not HEWL or myoglobin) eliminates the aggregation, especially at concentrations of 25 and 50 mg/mL (Figure 6A). FRAP experiments validate the micrograph observations (Figure 6B to E). The percent FRAP recovery of SB216763 is very low in buffer or in the presence of myoglobin crowders, but reaches close to 100% with the addition of 50 mg/mL BSA (see normalized fluorescence in Figure 6B). Also, D_confocal_ is much higher in the presence of 50 mg/mL BSA (Figure 6C). FRAP recovery is in line with the micrographs shown in Figure 6A. Titrating increasing concentrations of BSA into SB216763 solutions shows D_confocal_ values increasing to 15 µm^2^s^-1^ in the presence of 10 mg/mL BSA and to 25 µm^2^s^-1^ in the presence of 100 mg/mL BSA. These D_confocal_ values are consistent with solubilization of high molecular weight aggregates of SB216763. Interestingly, binding experiments of SB216763 to BSA, HEWL, and myoglobin showed affinities of 5.2±1.2 µM, 1.7±0.7 µM, and 7.3±1.8 µM, respectively (Figure 6 G-I, Table S1). Thus, while BSA solubilizes SB216763 much better than the other two proteins, we did not find a relation between solubilization of the small molecule and binding affinity.

**Figure 6:**
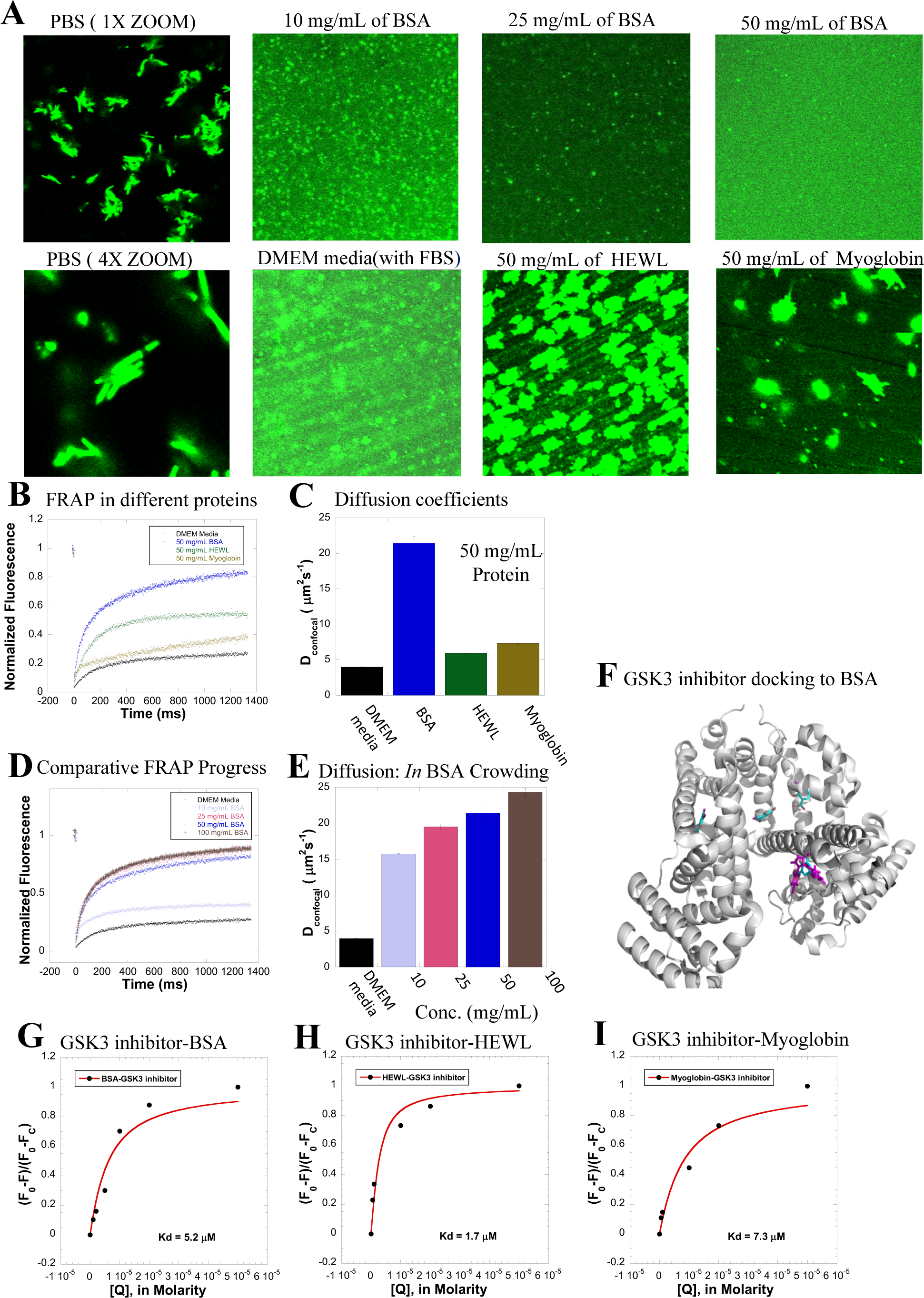
GSK3 inhibitor diffusion in protein crowders. (A) Confocal images of GSK3 inhibitor aggregates in PBS, DMEM media, BSA, HEWL and myoglobin. Note the disappearance of aggregation with increasing BSA concentration. (B) Averaged FRAP profiles (N=30; R=0.99 for each of the fits) and (C) diffusion coefficients for GSK3 inhibitor in the presence of 50 mg/mL BSA, HEWL and myoglobin. (D) Averaged FRAP profiles (N=30; R=0.99 for each of the fits) and (E) diffusion coefficients of GSK3 inhibitor in different concentrations of BSA protein. (F) BSA (grey) bound to 3,5-diiodosalicylic acid (PDB 4JK4, ligand with carbons in cyan). Docking of the GSK3 inhibitor (pink) to BSA (PDB 4F5S) was performed using different grid centers to capture each of the four binding sites for 3,5-diiodosalicylic acid. In one of the binding sites, the GSK3 inhibitor occupied a position similar to that of 3,5-diiodosalicylic acid, showing that the GSK3 inhibitor can bind to a rather buried binding site on BSA. (G-I): (F_0_-F)/(F_0_-F_c_) vs [Q] plots from fluorescence quenching experiments of GSK3 inhibitor-BSA (R= 0.97)/ HEWL (R= 0.98) /myoglobin (R= 0.98) systems in PBS buffer, where Q is the titrating drug concentration in molarity.

Visual inspection of the apo and holo crystal structures of the protein crowders shows that myoglobin does not have a crevice or buried binding site for small molecules larger than molecular oxygen, whereas HEWL and BSA do. Moreover, HEWL and BSA were shown to bind to small molecules and act as potential drug carriers (41, 46, 47). HEWL can accommodate small molecules in the crevice of its catalytic site, while BSA has four binding cavities that can accommodate small molecules (Figure 6F). Docking of the GSK3 inhibitor to BSA and to HEWL shows that the GSK3 inhibitor can bind to one of the cavities of BSA with a good docking score (-9.2 kcal/mol, Figure 6F) and to the catalytic site of HEWL with a less favorable docking score (-7.9 kcal/mol). The deep binding cavities in BSA that can accommodate the GSK3 inhibitor may explain why BSA is the only protein crowder that can reduce the aggregation of this small molecule. The diffusion coefficients computed from BD simulations did not show the trends obtained experimentally for the GSK3 inhibitor in the presence of protein crowders (Figure S1), indicating the importance of factors omitted in the BD model, such as small molecule aggregation and binding of small molecules within rather buried cavities in the protein crowder.

### Quinacrine Dihydrochloride

Quinacrine dihydrochloride (MW= 472.9 Da), whose chemical structure and spectroscopic properties are shown in Figure 1D, is among the first antimalarial drugs discovered. Recently, it was also suggested to have high potency against SARS-CoV-2 in an *in vitro* setting (48). Quinacrine does not display any significant aggregation issues in buffer solutions (Figure 7 A, B), and it is not adsorbed onto glass surfaces (Figure 3C). The addition of BSA, HEWL or myoglobin did not affect its diffusion coefficient (D_confocal_∼ 41- 43 µm^2^s^-1^, Figure 7 C, D). Control experiments in DMSO and in other media with and without the presence of Tween 20 did not alter the D_confocal_ values significantly, consistent with the observed solubility of quinacrine across multiple solutions (Figure 7A). Measuring the binding affinity of quinacrine to BSA, HEWL and myoglobin showed affinities of 4.6±1.1 µM, 4.7±0.8 µM and 3.0±0.4 µM, respectively (Figure 7 E-G, Table S1).

**Figure 7:**
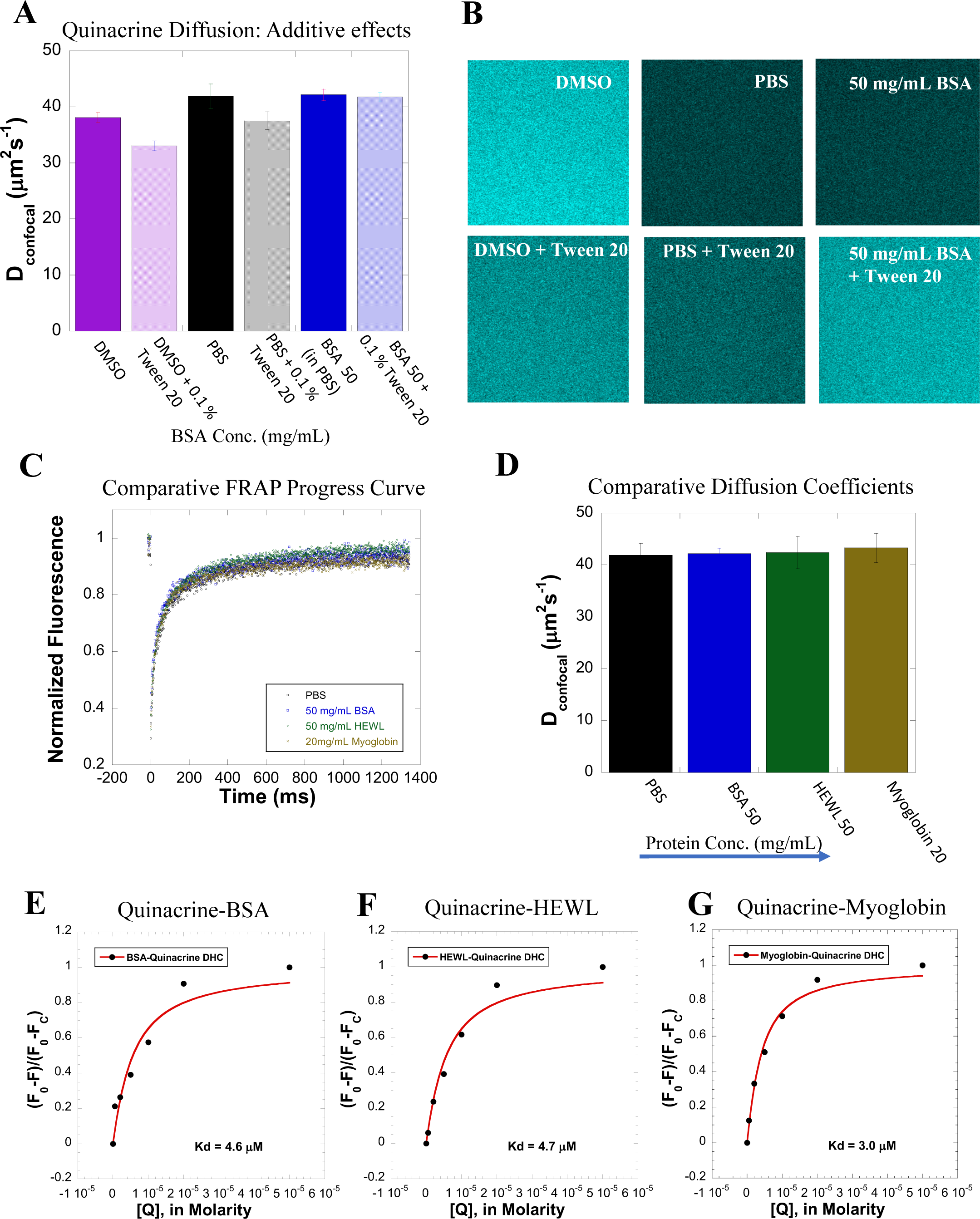
Quinacrine in protein crowders. Comparative (A) diffusion coefficients and (B) images from confocal microscope of Quinacrine DHC in diversified environments with or without presence of tween 20 are shown. (C) Comparative averaged FRAP profiles (R=0.99 for each of the fits) and (D) Comparative diffusion coefficients of Quinacrine DHC in presence of 50 mg/mL of BSA, HEWL, 20 mg/mL of Myoglobin and PBS buffer only are shown. (F_0_-F)/(F_0_-F) vs [Q] plots from fluorescence quenching experiments of Quinacrine DHC-BSA (R= 0.97)/ HEWL (R= 0.99)/Myoglobin (R= 0.99) system in PBS buffer is shown in (E-G), where Q is the titrating drug concentration in Molarity

As in the experiments, diffusion coefficients computed from BD simulations for quinacrine show no significant difference on adding 50 mg/mL protein crowder (Figure S1). One reason for the insensitivity to protein crowders is that quinacrine-protein contacts are less prominent compared to the other small molecules in BD simulations (Figure S3), as also shown by the absence of a peak in the quinacrine-HEWL RDF (Figure S4). However, the number of quinacrine-protein contacts was similar to that for the other small molecules for myoglobin and contacts were more pronounced for BSA. These differences are consistent with the larger positive charge on quinacrine than the other molecules (+2 e), resulting in greater solubility in aqueous solution and a net charge repulsion to HEWL and a net charge attraction to BSA, which however does not result in any significant difference in either the computed or the experimentally determined diffusion coefficients for quinacrine in 50 mg/mL BSA.

## Discussion

Studying the diffusion of low MW drugs in diverse environments is challenging, mostly because only a few drugs have the desired spectral properties. Nevertheless, studying their behavior in aqueous solution and in the presence of macromolecular crowdersis important as these environments are relevant for their administration and action. Here, we measured the diffusion of the four molecules in buffer solutions and solutions containing one of three different proteins as crowders, BSA, HEWL, and myoglobin. In addition, we determined the affinity of the small molecules for the three proteins using fluorescence quenching. The equilibrium dissociation constants are moderate (in the range of 1-100 µM). These moderate affinities explain the lack of co-elution in size exclusion chromatography (SEC) of the small molecules to the protein crowders (Figure S10, S11 and S12). Co-elution can be observed only for binding affinities better than 0.1 µM, which is not the case here. Yet, the addition of the proteins had a substantial effect on the diffusion of the small molecules. BD simulations and molecular docking revealed mechanistic details of the main experimental observations, providing further evidence for the different mechanisms of modulation of small molecule diffusion rates observed for the four compounds.

Each of the four small molecules studied behaved differently, providing excellent examples for different types of diffusional behavior of small molecules (Figure 8). Quinacrine dihydrochloride is a cationic charged, water-soluble compound. Its measured diffusion rate is not affected by the glass surface or the protein crowders. Fluorescein is a negatively charged, water-soluble compound. It forms interactions with proteins, which slow down its diffusion, mostly through its interaction with the surface adsorbed fraction of the proteins (Figures 3A, 3D and 4). The opposite is observed for DOX, a positively charged molecule, which diffuses slowly in PBS, but at a faster rate on the addition of protein crowders. A detailed analysis showed that DOX adsorbs to the glass surface and this is the reason for the slow diffusion. Last, SB216763 is a molecule with neutral net charge that is insoluble in water and forms large aggregates. However, addition of BSA as a carrier protein solubilizes SB216763, as observed from micrographs, the fraction of recovery, the increased diffusion rate, and its ability to bind in a rather buried binding site on BSA.

**Figure 8:**
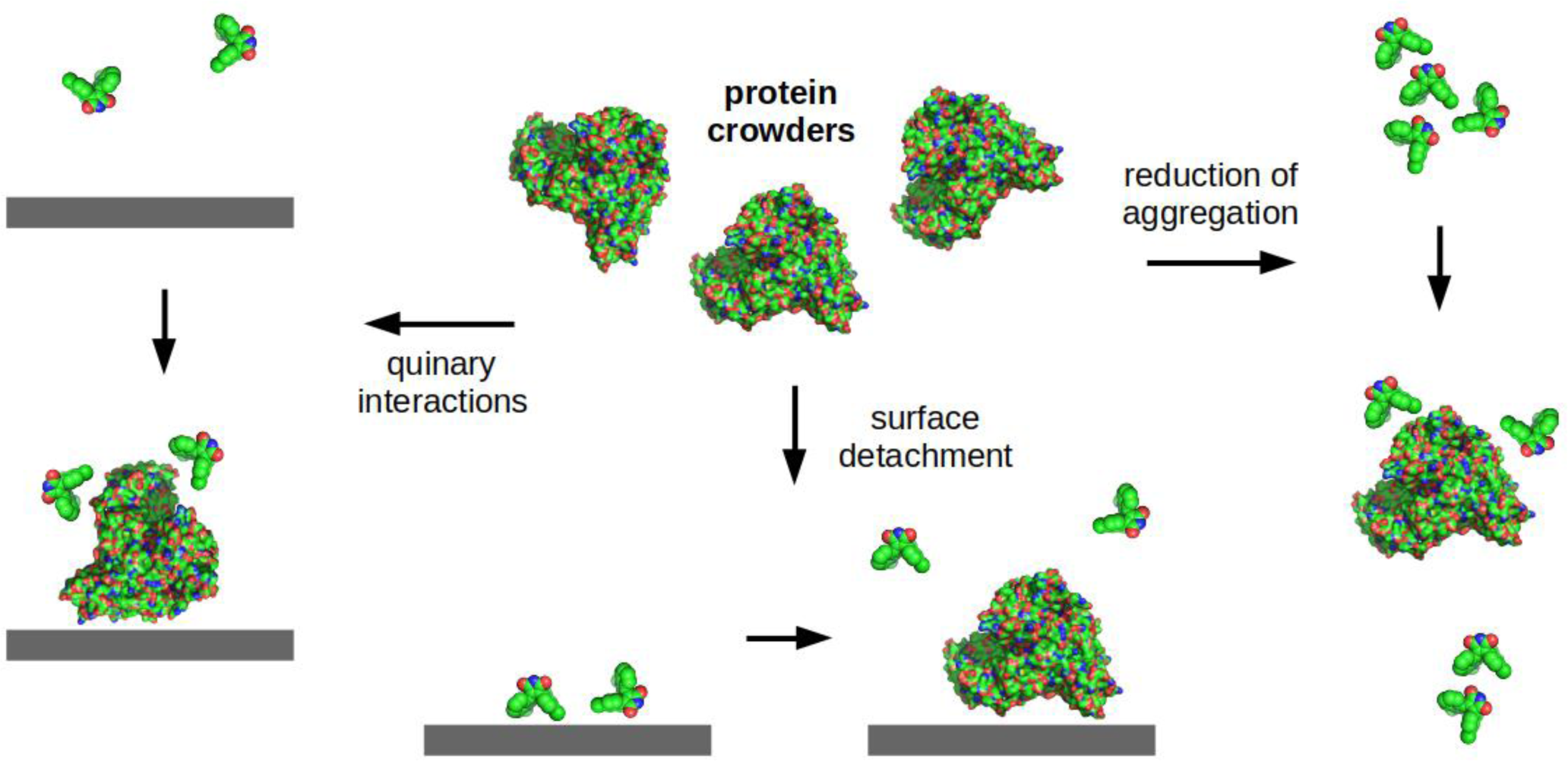
Effects of protein crowders on the diffusion of small molecules. The effects of protein crowders go beyond the slower diffusion due to excluded volume; the diffusion coefficients can also be slowed down due to quinary interactions or increased by surface detachment or reduced aggregation of the small molecules. The specific effect of the protein crowder on the diffusion coefficient of the small molecule depends on the physicochemical properties of the crowder and of the small molecule – which influence the protein-small molecule, protein-(glass) surface and small-molecule- surface interactions, as well as self-interactions.

A point of potential concern is the “slow” measured diffusion coefficients for the small molecules, which are much below those expected for these molecules in aqueous solution. A possible reason could be limitations in the Line-FRAP method, which may not allow the measurement of high diffusion coefficients (> 100 µm^2^s^-1^). If this would be a result of a “dead-time” between the end of the bleach and the first recovery data points, we should observe an incomplete bleach (as some recovery occurs before the first data measurement). However, the FRAP curves in Figures 2, 5, 6 and 7 do not show reduced bleach, even for the fastest diffusing molecules. This would suggest that for fast diffusing particles, the data analysis carried out on the experimental observations does not provide absolute diffusion coefficients. Therefore, we name the derived values D_confocal_. However, we consider the relative diffusion rates between the different conditions applied in this study to be robust, as the same data analysis procedure was carried out for all conditions. Unfortunately, methods such as FCS, which could provide an orthogonal approach for diffusion measurements, are not applicable to the small molecules used here, due to their low quantum yield.

In summary, by combining experiments and simulations, we observe that the effects of protein crowders on the diffusion of low molecular weight drug molecules go beyond the slowing of translational diffusion due to excluded volume, and find that the diffusion of drugs can also be slowed down due to quinary interactions, with the extent dependent on protein-surface interactions, or it can be increased by surface detachment or a reduction in aggregation of the small molecules (Figure 8).

## Materials and Methods

BSA, HEWL and myoglobin were purchased from Sigma-Aldrich. DOX was purchased from AdooQ Bioscience (catalog no. A14403). Fluorescein disodium salt was purchased from chemcruz (catalog no. sc-206026). The GSK3 inhibitor SB216763 (catalog no. ab120202), and quinacrine dihydrochloride (catalog no. ab120749), were purchased from Abcam. All other reagents used are described in a table format (Table S5, Key resources).

### Confocal microscopy and FRAP Analysis

Images were collected with an Olympus IX81 FluoView FV1000 Spectral/SIM Scanner confocal laser-scanning microscope (for details see Appendix). All image analyses were performed using FluoView software, and data analyses were performed using Kaleidagraph software version 4.1 (Synergy).

### Line-FRAP and Classical XY-FRAP

Line-FRAP was carried out in liquid drops as detailed in (36). For photobleaching, “Tornado” of 4 pixels (2x2) diameter was used in the simultaneous stimulus scanner. The unidirectional lines were scanned with time intervals of 1.256 ms 1000 times (equivalent to 1.256 s) in the majority of the measurements. We have used two simultaneous scanners during the FRAP experiments: one scanner (at 405 nm with the full intensity of 100%) for photobleaching and another scanner (at 440/515nm with weak intensity) for data acquisition. Fluorescence recovery plots were fitted to a double exponent growth curve. For further details see Appendix. Our previously developed Line FRAP protocol was employed for the final calculations of diffusion coefficients from the FRAP rates and averaged bleach sizes (36).

### Steady-State Fluorescence Quenching Assays

Steady-state fluorescence quenching experiments were carried out on a Tecan fluorescence plate reader instrument (for details see Appendix). The fluorescence emission spectra were recorded at λ_exc_ = 280 nm and λ_em_ from 300 to 450 nm, with the intensity at 344 nm (tryptophan) being used to calculate the dissociation constant (49). Fluorescence changes upon formation of a 1:1 complex is given by Eq. (2)

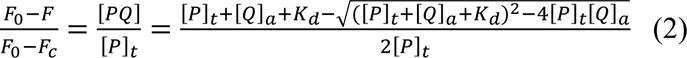

Where *F* is the measured fluorescence, *F0* is the starting fluorescence, *Fc* is the fluorescence of the fully complexed protein, *K*_d_ is the dissociation constant, [*P*]_t_ is the concentration of protein, and [*Q*]_a_ is the concentration of added ligand/quencher (small molecule/drug in our case). Data were fitted using Kaleidagraph version 4.1 (Synergy). The dissociation constants, *K*_d,_ are given in Table S1.

### Dynamic Light Scattering (DLS) Measurements

The hydrodynamic size (in nm) of the protein and protein/small molecule complexes in PBS buffer solution was measured using a Malvern’s Zetasizer Nano ZSP with a backscatter detection system at an angle of 173°. A minimum of three measurements were recorded for each sample. ZEN0040 disposable cuvettes with 400 μL of the sample were used. The equilibration time was about 15–30 min, and all the measurements were done at 25°C. The hydrodynamic size (in nm) was computed using Malvern’s Zetasizer software. Only auto-correlogram profiles (Correlation Function) with good quality fits were taken for final data consideration. Fluorescein:BSA concentration ratios were kept similar to the FRAP experimental conditions. In one measurement, the fluorescein concentration was increased up to 10 times (from 20µM to 200µM) that used in FRAP experiments to see if, at a higher concentration, fluorescein can alter the oligomerization state of BSA or not.

### Brownian dynamics simulations

BD simulations were performed using the Simulation of Diffusional Association (SDA) 7 software package (50), version 7.2, available at https://mcm.h-its.org/sda. BD simulations were performed using a time step of 0.5 ps, ionic strength of 190 mM and periodic boundary conditions. Small molecules and proteins were kept rigid, and the same conformations of each small molecule and protein were used for all simulations. The detailed setup is described in the Appendix.

First, all simulated systems were subjected to a 0.2 μs-length BD simulation with interactions between molecules modelled by only a soft-core repulsion energy term in order to resolve any steric clashes in the initial simulation box generated by the SDA tool genbox. After this, BD production simulations were performed for 10 μs. For all BD simulations, radial distribution functions and the number of small molecule-protein or small molecule- small molecule contact interactions were computed using SDA7. Contacts were defined as present when heavy atoms (at least one in each molecule) were within a distance of 4.5 Å of each other. Snapshots were analysed at intervals of 500 picoseconds and values averaged over the last 10 microseconds of the simulations. Computation of diffusion coefficients for small molecules and their docking to proteins is described in the Appendix.

## Acknowledgments

We gratefully acknowledge Stefan Richter (HITS) for technical support of the computational work, Neil J. Bruce (HITS), Daria B. Kokh (HITS) and Huan-Xiang Zhou (<u>University of Illinois at Chicago</u>) for helpful discussions, and Gaurav Ganotra (HITS) for help with the parameterization of small molecules. This work was supported by the Israel Science Foundation grant no. 1268/18 (GS), a Capes-Humboldt postdoctoral scholarship to AN-A (Capes process number 88881.162167/2017-01), funds from the Cluster of Excellence CellNetworks (DFG, EXC81) to AN-A, the European Union’s Horizon 2020 Framework Programme for Research and Innovation under Grant Agreements 785907 and 945539 (Human Brain Project SGA2 and SGA3) (RCW), and the Klaus Tschira Foundation (AN-A, RCW).

## Authors Contributions

DD carried out the experiments. AA performed the computer simulations. DD, AA, RCW and GS conceived the study, analyzed the data and wrote the manuscript.

## Supporting information

### Supplemental Methods

#### Confocal microscopy and FRAP Analysis

Images were collected with an Olympus IX81 FluoView FV1000 Spectral/SIM Scanner confocal laser-scanning microscope, using 60X DIC oil-immersion objective, N.A. 1.35. For fluorescein disodium salt fluorescence measurements, excitation was done at 440 nm, using a diode laser at an output power of 1-4 % of maximal intensity for high to low concentrations, whereas emission was recorded from 520 to 550 nm using the spectral detection system. For doxorubicin, excitation was done at 515 nm diode laser using 5-10% of the maximal intensity, and emission were collected from 540-640 nm. For quinacrine di hydro chloride and GSK3 inhibitor, excitation was done at 488 nm laser using 1-2% of the maximal intensity, while emission was collected from 502-560 nm with SDM560 emission dichromator cut off filter.

#### Line-FRAP and Classical XY-FRAP

Line-FRAP was carried out in liquid drops. For photobleaching, “Tornado” of 4 pixels (2x2) diameter was used in the simultaneous stimulus scanner. This is the smallest area achievable using Tornado. The circle area of the bleach was kept precisely in the middle of the scanning line. The unidirectional lines were scanned with time intervals of 1.256 ms 1000 times (equivalent to 1.256 s) in the majority of the measurements. The number of scans before, during, and after photobleaching was 10, 42, and 948, respectively. Photobleaching was achieved by the simultaneous laser at 405-nm excitations with 63 millisecond durations, using at full intensity (100%). The simultaneous scanner moved at a speed of 100 µs/pixel to perform an efficient photobleach. We have used two simultaneous scanners during the FRAP experiments: one scanner (at 405 nm with the full intensity of 100%) for photobleaching and another scanner (at 440/515nm with weak intensity) for data acquisition. For all the drugs, bleach was performed by 405 nm laser, whereas for main excitations, Fluorescein 440 nm laser (1-4%); GSK3 inhibitor 440 nm laser (0-1%) were used. Emission collections were done from 520-550 nm for Quinacrine DHC and GSK3 inhibitor. Using the Olympus IX81 FluoView FV1000 Spectral/SIM Scanner confocal laser-scanning microscope, using Tornado (which requires SIM scanner to be loaded) greatly enhances bleaching efficiency. In addition, it shortens the time to obtain the first measurement after bleach (which is immediate in this mode). This property is highly beneficial for Line-FRAP measurements, where the time scale of data acquisition plays an important role. The Fluoview SIM scanner unit synchronizes laser light simulation with confocal and multiphoton imaging to avoid interruption to image observation during laser stimulation or manipulation. We have varied the intensity of the lasers to achieve a good signal/noise ratio. Fluorescence recovery plots were fitted to a double exponent growth curve. FRAP experiments were also performed inside the PBS buffer drops and in crowding conditions. Glass plate dish containing cover slips are used for microscopic measurements.

#### Steady-State Fluorescence Quenching Assays

Steady-state fluorescence quenching experiments were carried out on a Tecan fluorescence plate reader instrument. BSA, Myoglobin, and HEWL solutions of 2 µM strength in PBS 1X (pH=7.4) were prepared. Different small molecules are added in the protein solutions, maintaining the final concentrations between 1 to 100 µM. The final volume of each mixture was strictly 200 µL. The whole set of experiments was performed in a 96-well microplate system (black, flat bottom, Fluotrac).

#### Brownian dynamics simulations

The initial coordinates for HEWL, BSA and myoglobin were obtained from the PDB files 1HEL (4), 4F5S (5) and 1DWR (6), respectively. Partial atomic charges and protonation states for the protein crowders at pH 7.2 were computed using pdb2pqr (7, 8). Partial atomic charges for the heme group of myoglobin were obtained from previous work (9).

The initial coordinates for the small molecules were obtained from conversion of SMILES to PDB format using Babel (10). The protonation state of the molecules at pH 7.2 was computed using Epik (11) in Maestro (Schrödinger, LLC, New York, NY). The log P and number of violations to Lipinski’s Rule of Five were computed using QikProp in Maestro (Schrödinger, LLC, New York, NY). The structures of the four small molecules were energy minimized using Maestro and then submitted to quantum mechanical calculations in GAMESS (12) using HF and the 6-31G** basis set to obtain RESP (13, 14) partial atomic charges.

BD simulations were performed using the Simulation of Diffusional Association (SDA) 7 software package (15), version 7.2, available at https://mcm.h-its.org/sda. All simulation boxes had the same composition: 80 small molecules of one type (fluorescein, doxorubicin, quinacrine or GSK3 inhibitor), or 80 small molecules and 440 protein crowders of one type (BSA, HEWL or myoglobin), resulting in a crowder:small molecule ratio of 5.5, which mimics the experimental conditions. Different crowder concentrations were achieved by changing the size of the cubic periodic simulation box (Table S6).

Three replica BD simulations were performed for each system. BD simulations were performed using a time step of 0.5 ps, ionic strength of 190 mM and periodic boundary conditions. Small molecules and proteins were kept rigid, and the same conformations of each small molecule and protein were used for all simulations. The translational and rotational diffusion coefficients of the small molecules and proteins at infinite dilution in aqueous solution, necessary to perform BD simulations, were computed from HYDROPRO (16), with a radius of the atomic element (AER) of 2.9 Å for proteins and 1.2 Å for small molecules (see details below, Table S7). Hydrodynamic interactions were computed using a mean-field model (17). The Stokes radii of the small molecules and proteins, necessary to compute hydrodynamic interactions, were calculated from the solvent-accessible volume estimated from a single point Poisson-Boltzmann calculation performed using AMBER 2016 (18) (Table S2). A radius of four times the Stokes radius of the protein was used to define the local volume for computing hydrodynamic interactions. The forces between molecules were modelled by computing electrostatic interaction, electrostatic desolvation and non-polar desolvation terms from the interactions between the atoms of each molecule and precomputed potential grids on the other molecules. Effective charges computed for the proteins (19) and for the small molecules (20) were used to calculate electrostatic interactions during the BD simulations. The grid spacing for protein crowders and small molecules was 0.75 Å for simulations with doxorubicin and 0.65 Å for simulations with the other small molecules. The lengths of the sides of the cubic grids for the small molecules were 97 grid points (electrostatic potential) and 80 grid points (for the other, shorter-range potentials). The lengths of the sides of the cubic grids were 161 grid points (electrostatic potential) and 135 grid points (for the other potentials) for HEWL, 225 grid points (electrostatic potential) and 193 grid points (for the other potentials) for BSA, and 225 grid points (electrostatic potential) and 200 grid points (for the other potentials) for myoglobin.

First, all simulated systems were subjected to a 0.2 μs-length BD simulation with interactions between molecules modelled by only a soft-core repulsion energy term in order to resolve any steric clashes in the initial simulation box generated by the SDA tool genbox. After this, BD production simulations were performed for 10 μs.

The parameters and grids described above were also employed to perform BD simulations in the presence of a silica surface to mimic glass. The parameters for the silica surface were obtained from a previous BD simulation study (21), and a similar simulation setup was used here. The silica surface was represented as a homogeneously charged graphite lattice surface with a charge density of -0.0013 e/ Å ^2^. The surface was assigned a charge distribution of -0.0032 e/atom. The x and y dimensions of the simulation box and of the silica surface were 340 Å. The desired crowder concentration was achieved by changing the height (z axis) of the simulation box. In the starting configuration for BD simulations, a buffer region of 120 Å separated the surface and the solutes. Before inclusion of the surface and buffer region in the system, systems were subjected to an initial 0.2 μs-length BD simulation with interactions between molecules modelled by only a soft-core repulsion energy term in order to resolve any steric clashes in the initial simulation box. After this, BD production simulations were performed for 10 μs.

In the analysis of molecule adsorption, a molecule was considered adsorbed when its distance from the surface was less than 4 times the Stokes radius of the molecule, as done previously (21).

For all BD simulations, radial distribution functions and the number of small molecule-protein or small molecule-small molecule contact interactions were computed using SDA7. Contacts were defined as present when heavy atoms (at least one in each molecule) were within a distance of 4.5 Å of each other. Snapshots were analysed at intervals of 500 picoseconds and values averaged over the last 10 microseconds of the simulations.

### Computation of diffusion coefficients for small molecules

HYDROPRO (16) was parameterized to reproduce the experimental translational and rotational diffusion coefficients of proteins. One parameter, the radius of the atomic elements (AER), was adjusted to reproduce the experimental results. A procedure similar to that followed in the original publication (16) was followed to obtain an AER value for small molecules:

1. estimate translational diffusion coefficients for small molecules using varying values of AER (1, 2, 3 and 4 Å);
2. do a linear fit for the graph AER values vs. translational diffusion coefficients for each small molecule, and calculate the AER value that reproduces the experimental translational diffusion coefficient;
3. calculate the final AER value as the average of the AER values obtained for each small molecule.

Following the procedure above, an AER value of 1.2 Å was obtained for small molecules. Table S7 shows the computed diffusion coefficients for molecules with known experimental diffusion coefficients using AER values before and after the reparameterization for small molecules.

### Docking of the small molecules to the proteins

Docking was performed using Autodock Vina (22), a cubic grid with a spacing of 0.375 Å and 70 grid points, and default parameters. The grid was centered on the catalytic site of HEWL (PDB 1HEL) or on one of the four binding cavities of BSA (PDB 4F5S), identified by the presence of 3,5-diiodosalicylic acid in the complex with BSA in PDB 4JK4 (23). The docking pose with the highest score was retained.

---

**Figure S1.**
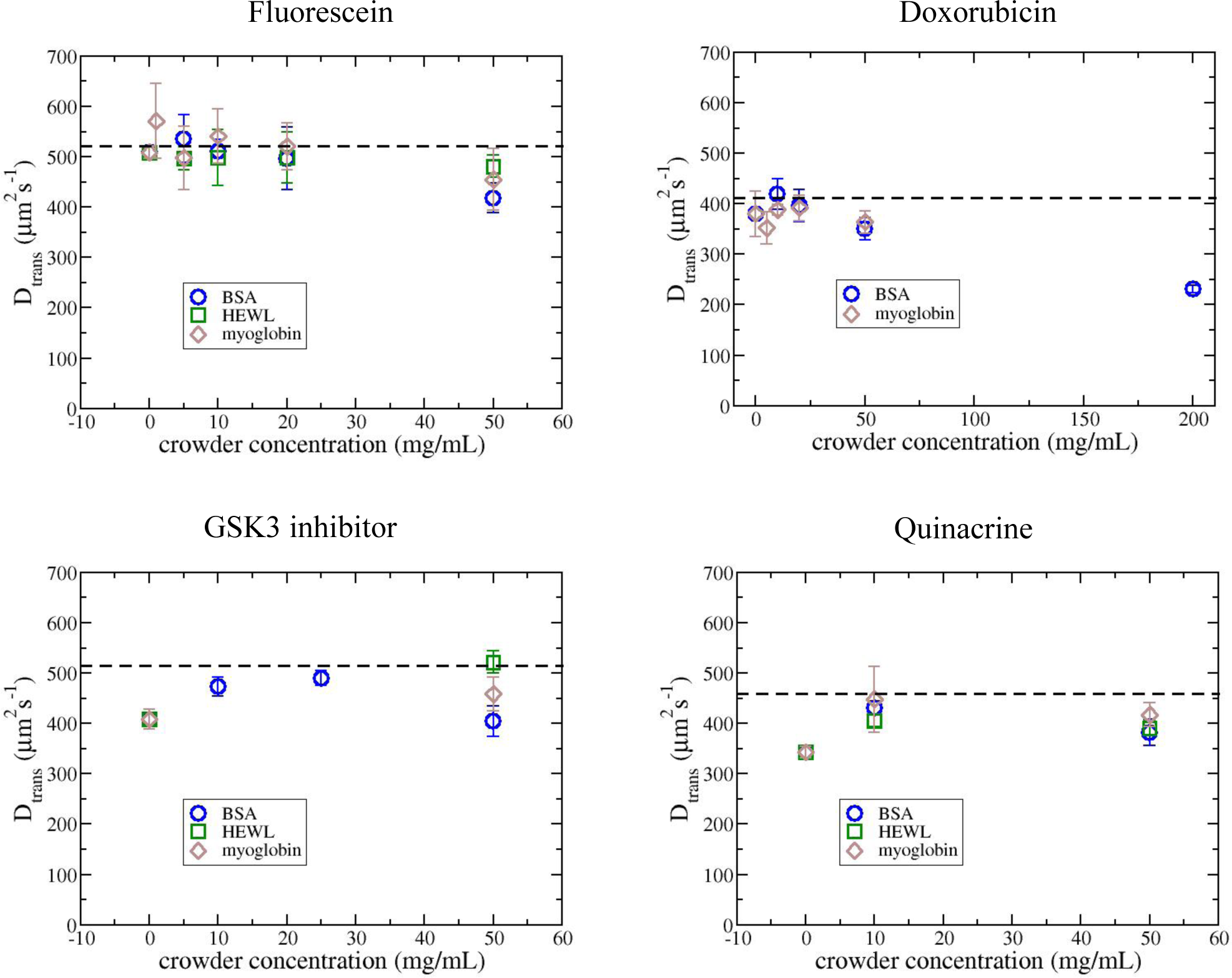
Translational diffusion coefficients of the four small molecules in the presence of protein crowders, obtained from BD simulations: The results show that the effects of the crowders modelled in these simulations (excluded volume, electrostatic and hydrophobic interactions between solutes) result in small or no reductions in the computed diffusion coefficients of the small molecules at crowder concentrations up to 50 mg/mL. As expected, a significant reduction in diffusion coefficient is seen for doxorubicin at a BSA crowder concentration of 200 mg/mL. Each point is the average ± standard deviation of 3 replica BD simulations, each of 10 microseconds duration. Horizontal dashed black line: translational diffusion coefficients of the small molecule in the absence of crowder (infinite dilution) computed using HYDROPRO. The systems were composed of 440 protein crowders and 80 small molecules.

**Figure S2.**
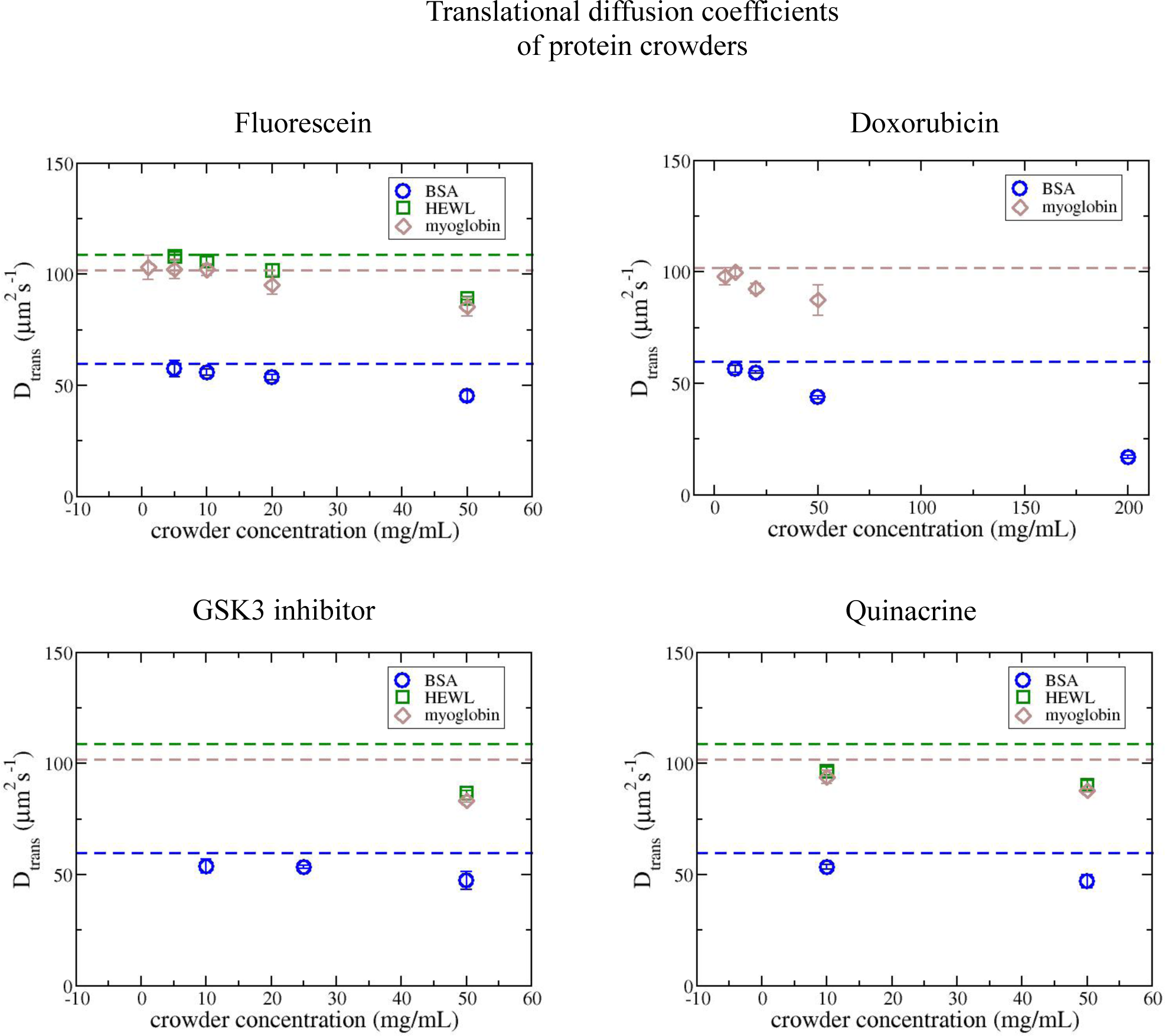
Translational diffusion coefficients of protein crowders obtained from BD simulations performed in the presence of small molecules: The results show that the effects of the crowders modelled in these simulations (excluded volume, electrostatic and hydrophobic interactions between solutes) result in modest reductions (up to 20%) in the computed diffusion coefficients of the crowders at crowder concentrations up to 50 mg/mL. As expected, a significant reduction in diffusion coefficient is seen for BSA at a BSA crowder concentration of 200 mg/mL in simulations with doxorubicin. Each point is the average ± standard deviation of 3 replica BD simulations, each of 10 microseconds duration. Horizontal dashed lines: translational diffusion coefficients of the protein crowders in infinite dilution computed using HYDROPRO. The systems were composed of 440 protein crowders and 80 small molecules.

**Figure S3.**
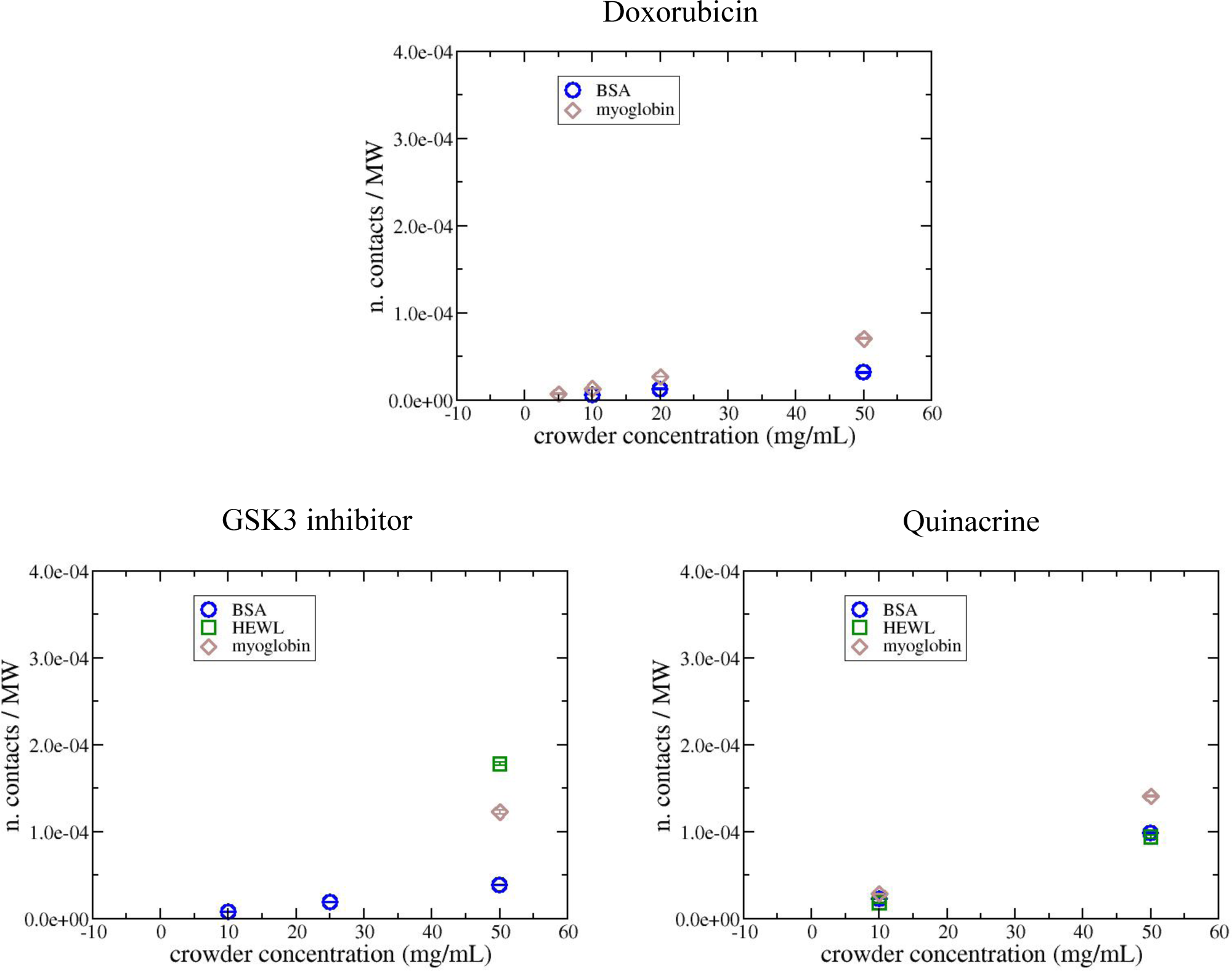
Ratios of the number of protein-small molecule contact interactions/protein molecular weight in the BD simulations of small molecules and protein crowders. The ratio is computed as the number of protein-small molecule contact interactions in BD simulations with different protein crowders divided by the molecular weight (MW) of the protein crowder (66637, 17820 and 14331 g/mol for BSA, myoglobin and HEWL, respectively). The number of contacts increases with increasing crowder concentration in a protein crowder and small molecule dependent fashion. Each point is the average ± standard deviation from 3 replica BD simulations, each of 10 microseconds duration. The corresponding plot for fluorescein is shown in Figure 4 of the main text.

**Figure S4.**
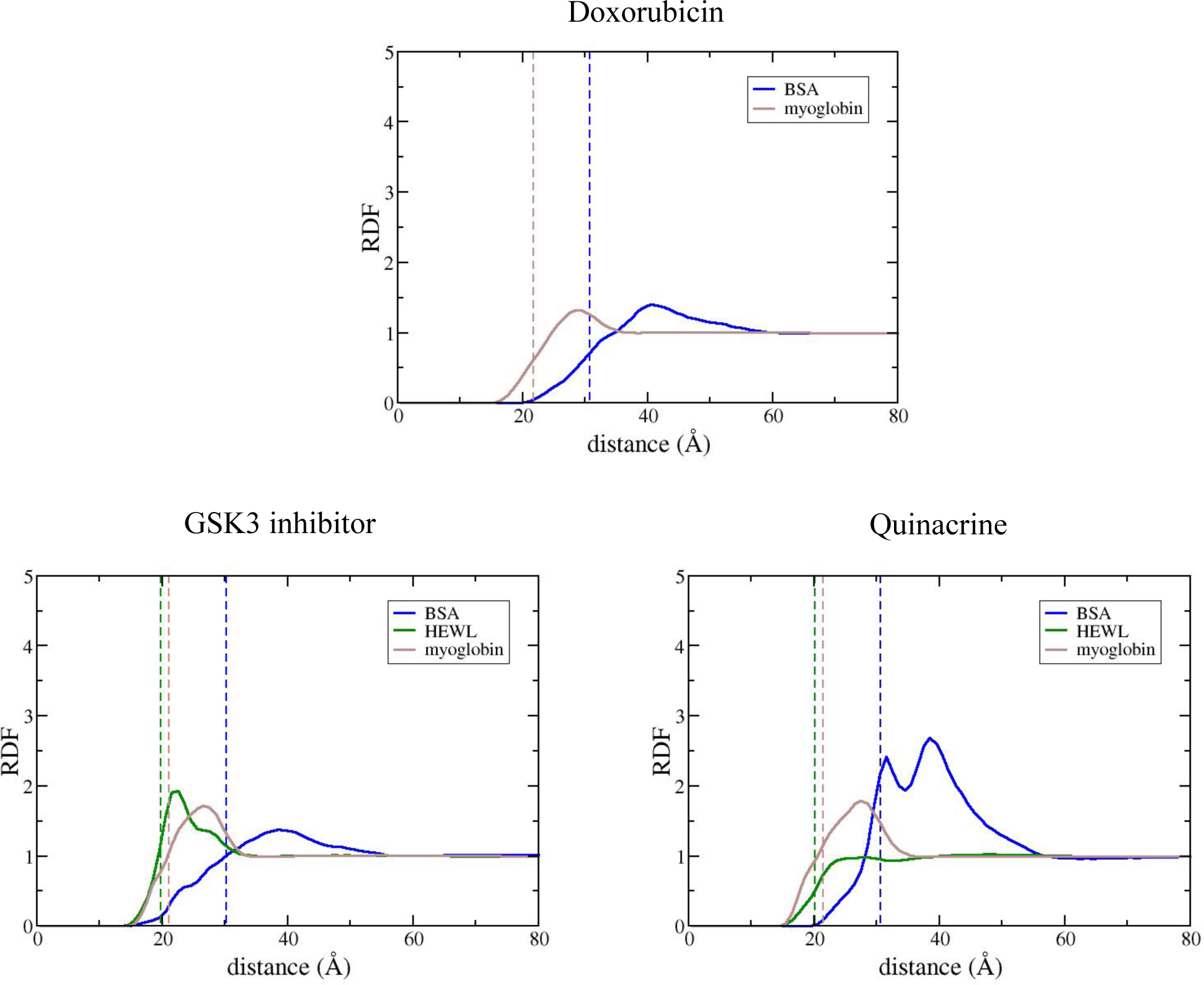
Radial distribution functions for protein-small molecule distances. RDFs for protein-small molecule distances observed in BD simulations with different protein crowders showing differences in the interactions of the three drugs with the three protein crowders. Each curve was computed from one BD simulation of 10 microseconds duration performed with 50 mg/mL of crowder (pH 7.2, ionic strength of 190 mM). The dashed lines indicate the sum of the Stokes radii of the respective protein crowder and small molecule. The Stokes radii of the proteins are 25.7 Å, 16.6 Å and 15.3 Å for BSA, myoglobin and HEWL, respectively, and the Stokes radii of the small molecules are 5.1 Å, 4.5 Å and 4.9 Å for doxorubicin, GSK3 and quinacrine, respectively. The corresponding plot for fluorescein is shown in Figure 4 in the main text.

**Figure S5.**
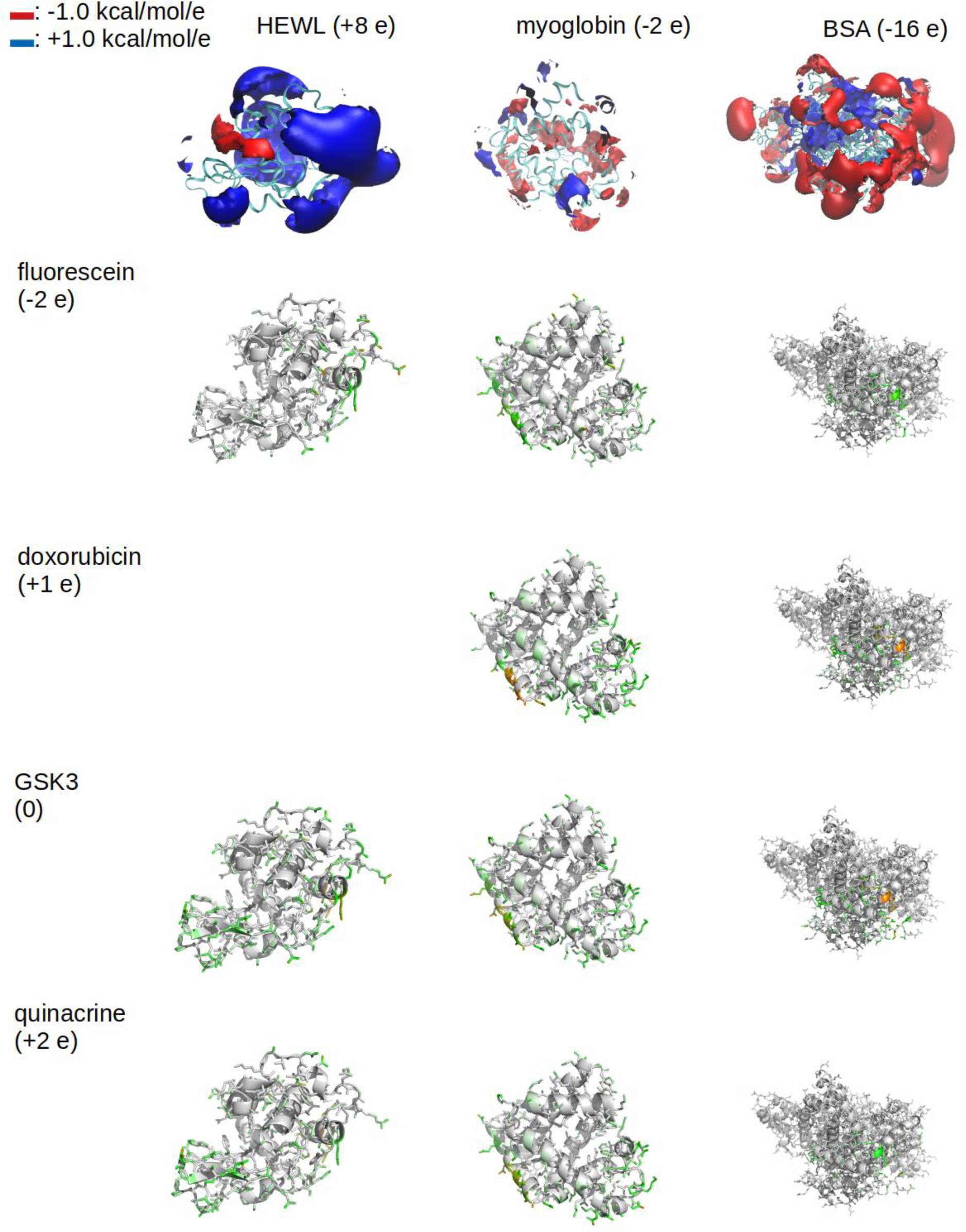
The four small molecules interact with similar regions of the same protein crowder in BD simulations. First row: molecular electrostatic potential isocontours for the three protein crowders (pH 7.2, ionic strength of 190 mM). Remaining rows: atoms with higher probability of making protein-small molecule interactions are shown in brighter colors (scale: gray (low probability of interaction) – green – orange (high probability of interaction)). Probabilities are computed for each system from one BD simulation of 10 microseconds duration with 50 mg/mL of protein crowder (pH 7.2, ionic strength of 190 mM).

**Figure S6:**
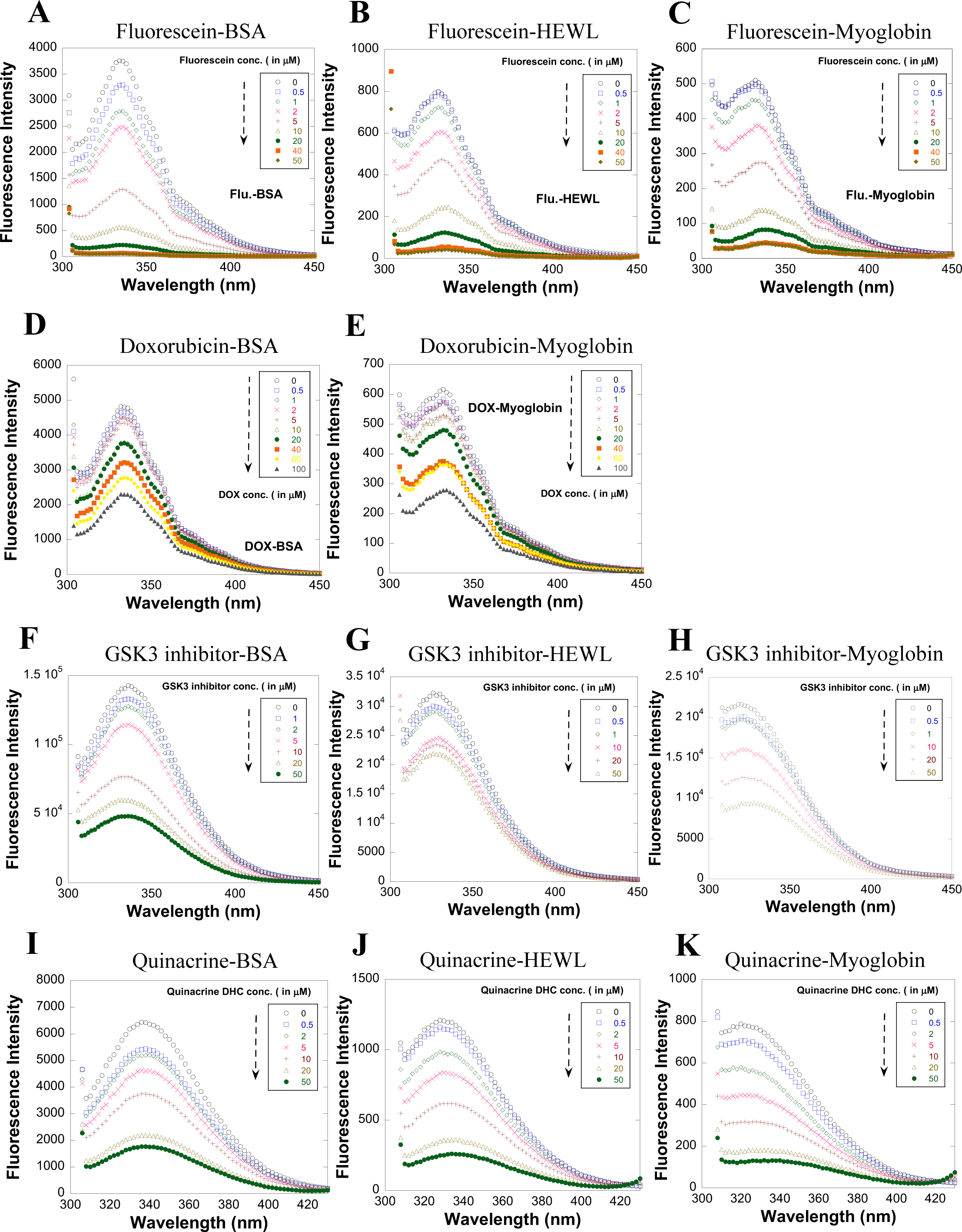
Steady-state fluorescence quenching assay: Fluorescence emission spectra of drug-BSA, drug-myoglobin, drug- HEWL systems in PBS buffer at pH 7.4 and 25°C presented for (A) Fluorescein-BSA, (B) Fluorescein-HEWL, (C) Fluorescein-myoglobin, (D) DOX-BSA, (E) DOX-myoglobin, (F) GSK3 inhibitor-BSA, (G) GSK3 inhibitor-HEWL, (H) GSK3 inhibitor-myoglobin, (I) Quinacrine DHC-BSA, (J) Quinacrine DHC-HEWL, and (K) Quinacrine DHC-myoglobin. Protein concentrations were fixed at 2 µM for titration. The small molecule drug concentrations used (in µM) are shown in the inset legends with the dashed downwards-pointing arrow indicating increasing concentration.

**Figure S7:**
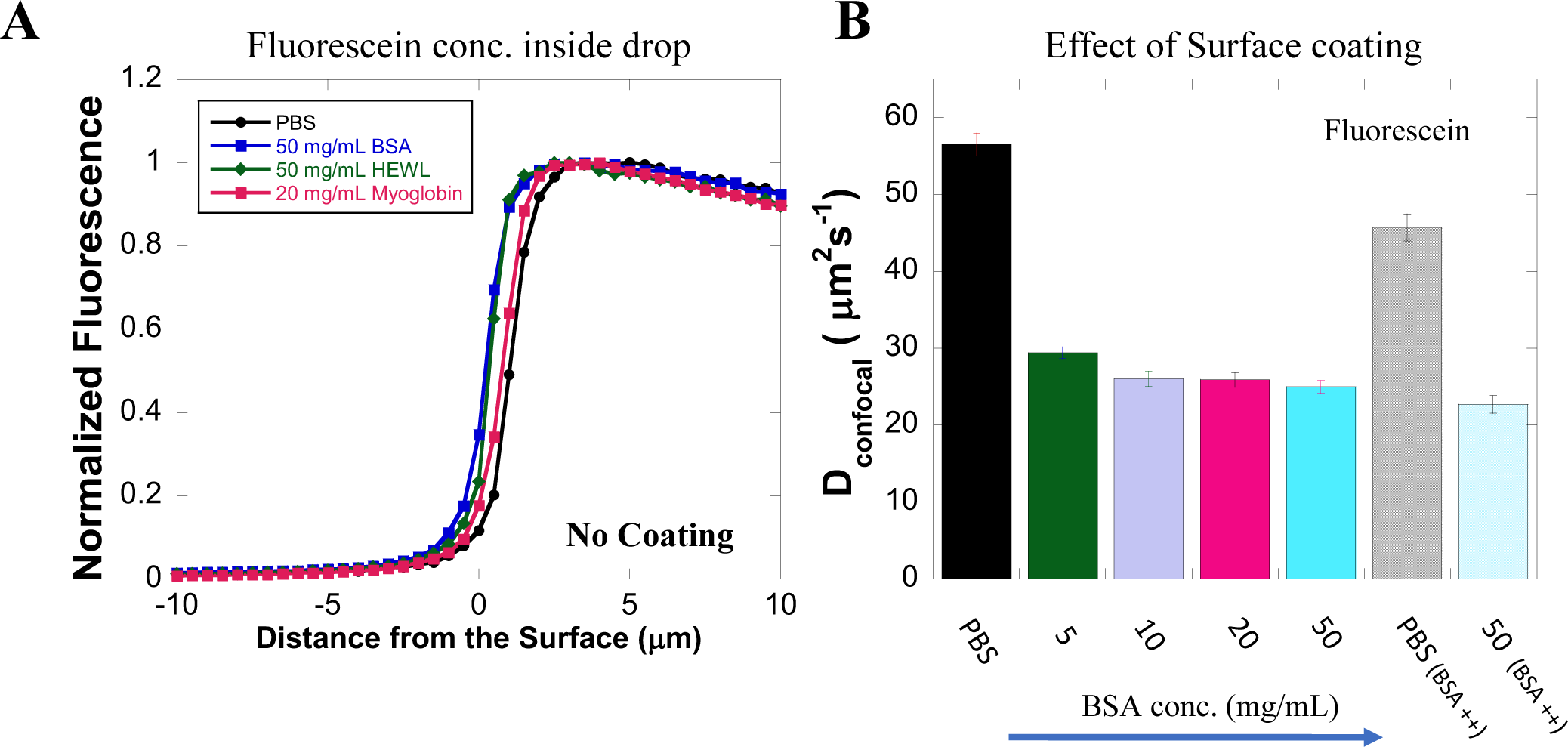
Adsorption of fluorescein to a glass surface in the absence and presence of protein crowders. (A) Determination of drop homogeneity of fluorescein in different crowding environments and in PBS buffer solution without surface coating. (B) Effect of surface coating by BSA on the diffusion coefficient of fluorescein in the absence and presence of BSA protein crowders. Prior coating of the glass surface with BSA results in small reductions in the respective D_confocal_ values in PBS and 50 mg/mL BSA. Here ending with ++ symbol denotes the glass coating.

**Figure S8:**
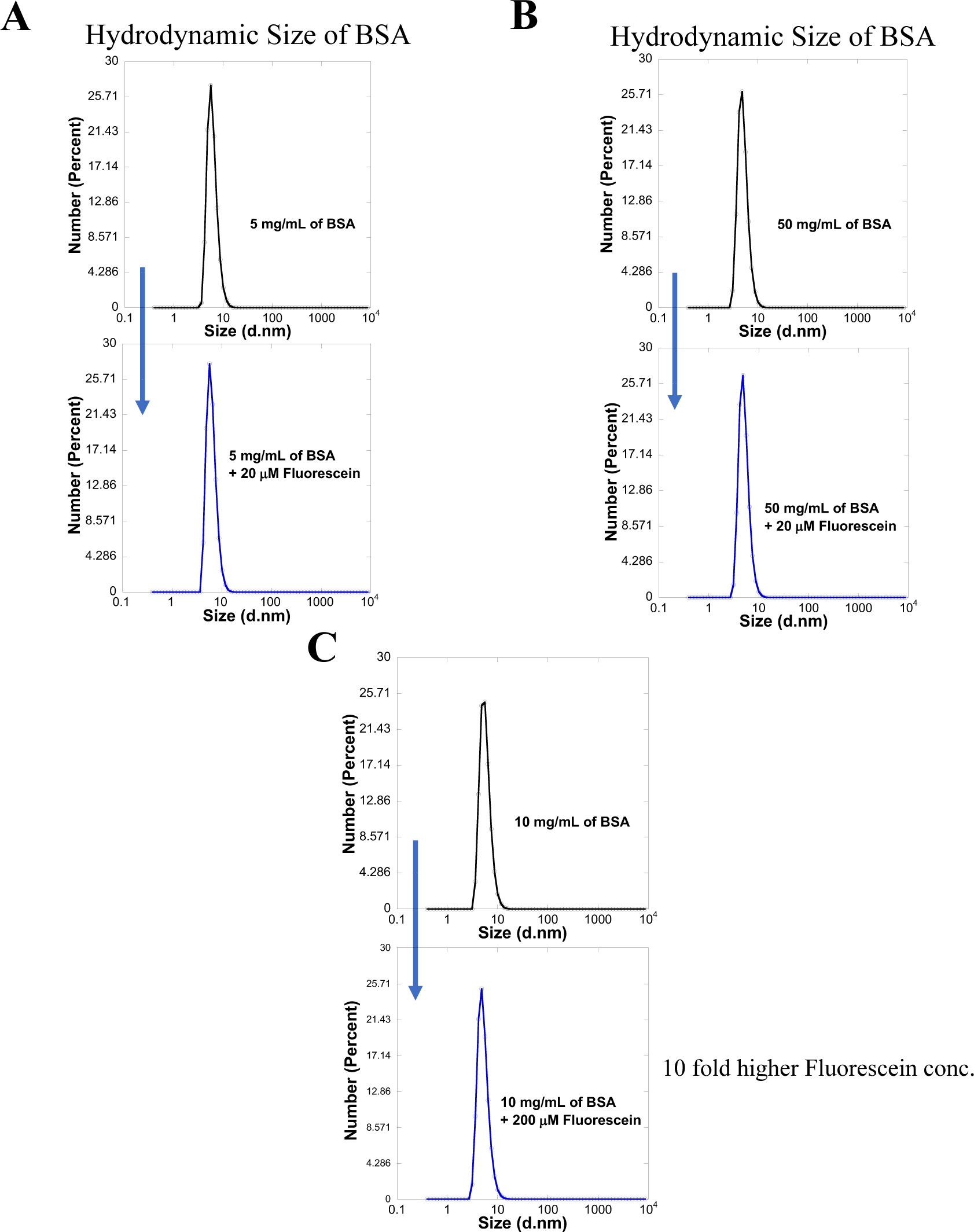
Hydrodynamic size measurements by dynamic light scattering of BSA and BSA/fluorescein mixtures: (A-C) The hydrodynamic size (in nm) of BSA in the absence and presence of fluorescein at different BSA concentrations is shown. The addition of fluorescein did not result in a significant change in the hydrodynamic radius of the protein (even when added at a concentration 10 times higher than used in FRAP experiments (C), indicating that the interaction of fluorescein with BSA does not result in significant protein oligomerization.

**Figure S9:**
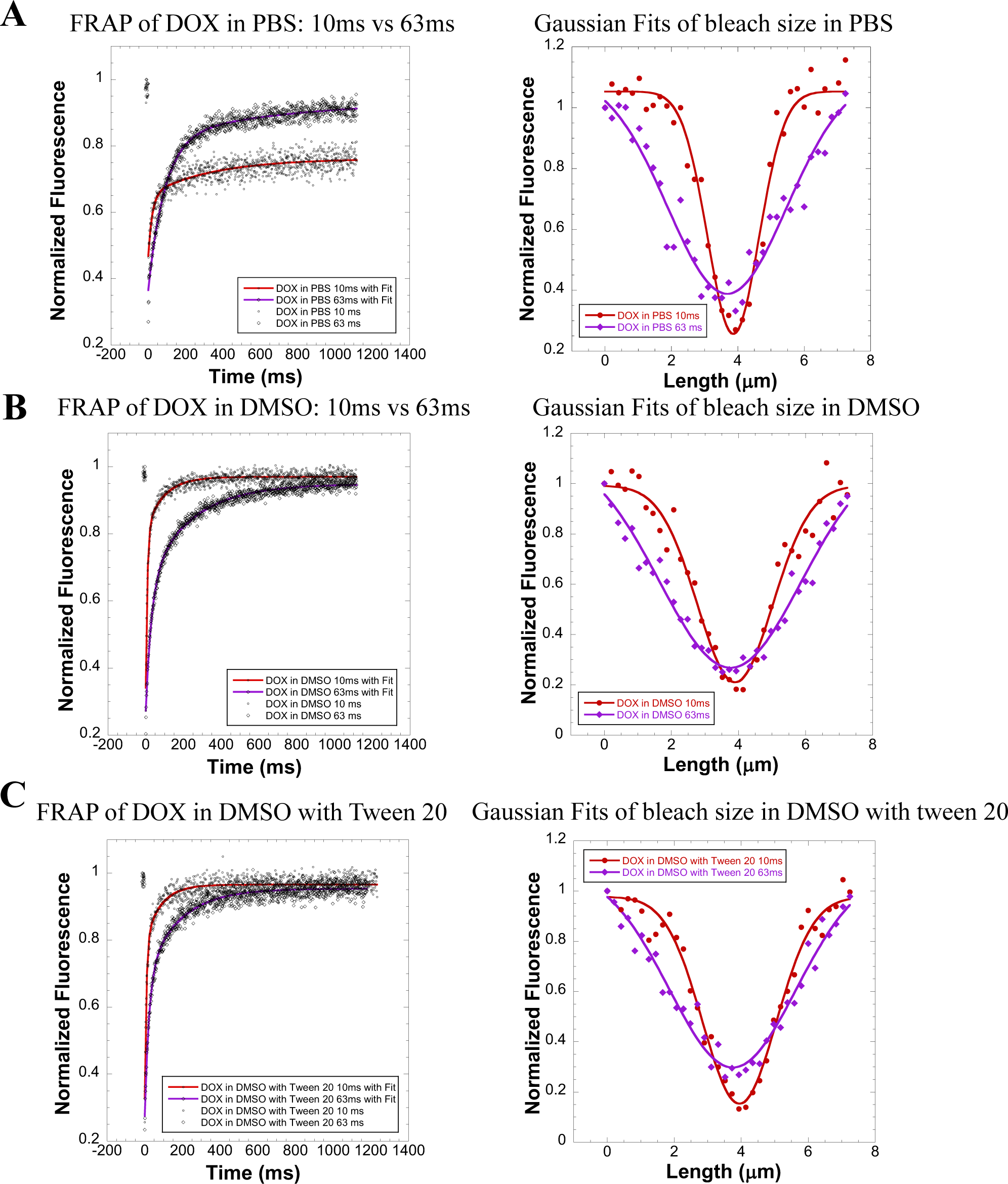
10 ms vs 63 ms bleach pulse comparison: Comparison of normalized FRAP recovery profiles and the corresponding Gaussian bleach size profiles with fits for 10 ms vs 63 ms bleach pulses, where doxorubicin is in (A) PBS buffer, (B) pure DMSO, and (C) DMSO with Tween 20.

**Figure S10:**
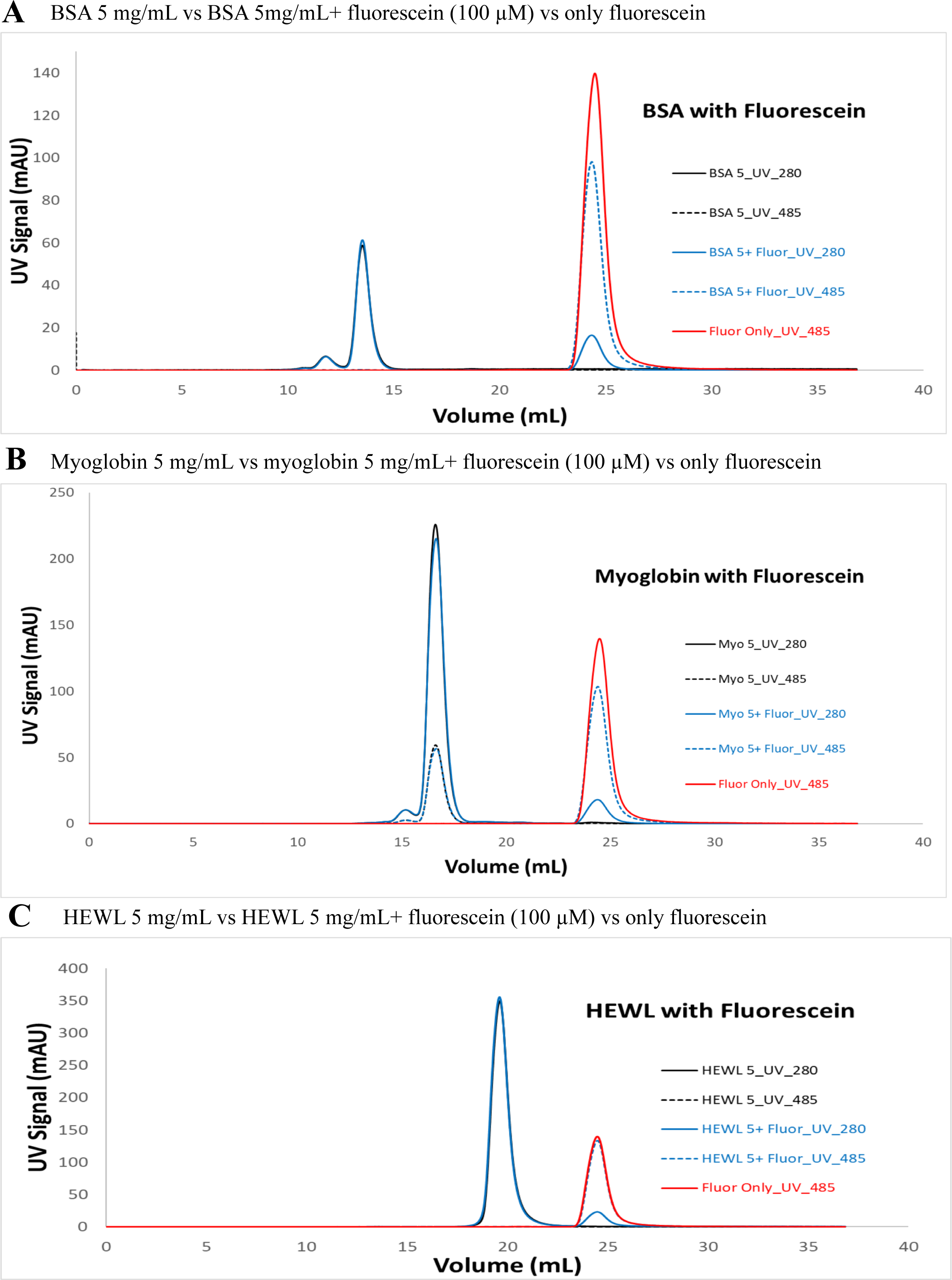
Size exclusion chromatography profiles of proteins and protein-bound fluorescein: Comparative size exclusion chromatography profiles are shown of (A) BSA, BSA-bound fluorescein and only fluorescein in PBS buffer solution, (B) Myoglobin, myoglobin-bound fluorescein and only fluorescein in PBS buffer solution, and (C) HEWL, HEWL-bound fluorescein and only fluorescein in PBS buffer. Simultaneous measurements were done at 280 nm and 485 nm wavelengths.

**Figure S11:**
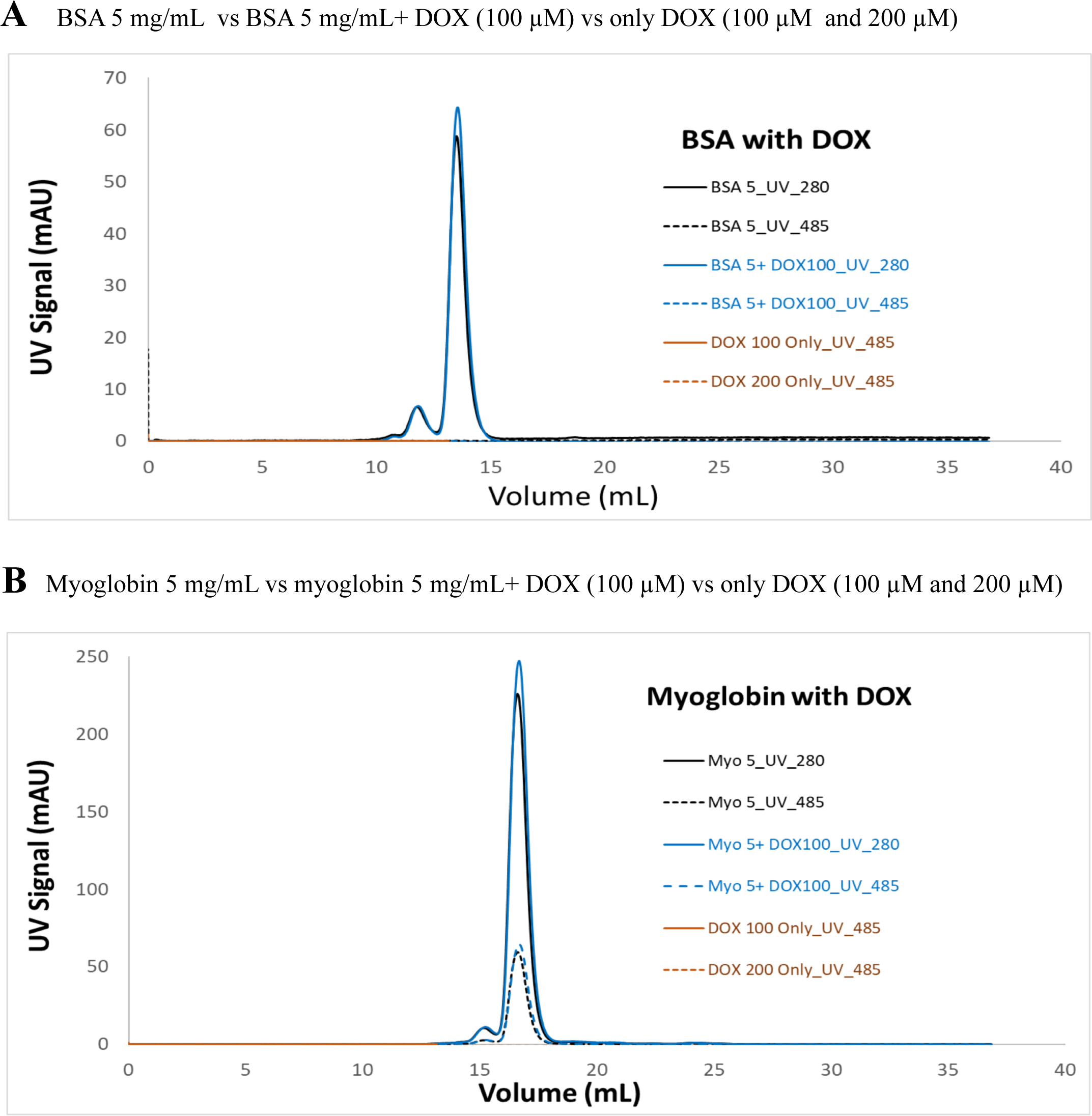
Size exclusion chromatography profiles of proteins and protein-bound DOX: Comparative size exclusion chromatography profiles are shown of (A) BSA, BSA-bound DOX and only DOX in PBS buffer solution, (B) Myoglobin, myoglobin-bound DOX and only DOX in PBS buffer solution. Simultaneous measurements were done at 280 nm and 485 nm wavelengths.

**Figure S12:**
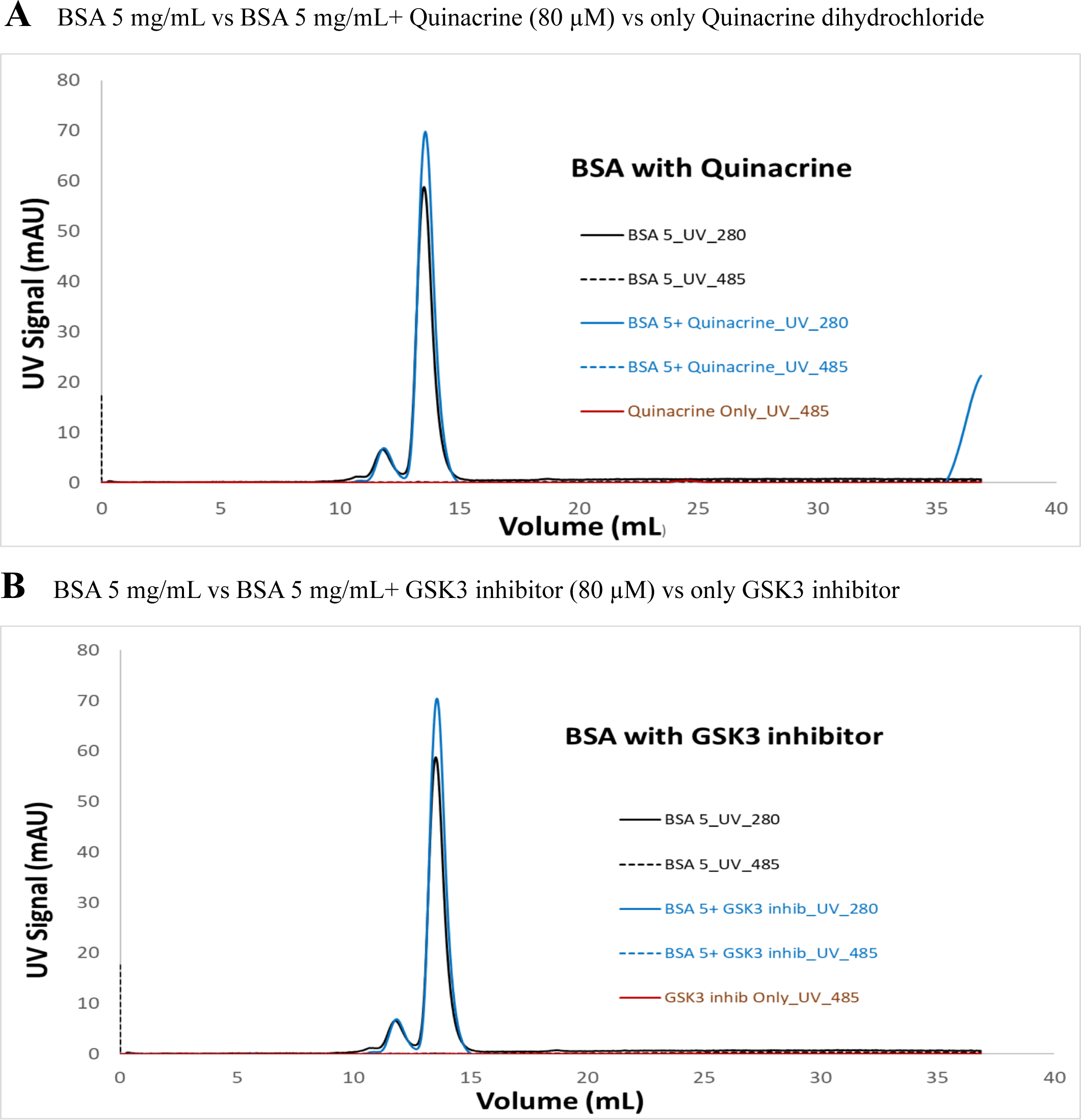
Size exclusion chromatography profiles of proteins and protein-bound quinacrine and GSK3 inhibitor: Comparative size exclusion chromatography profiles are shown of (A) BSA, BSA-bound quinacrine and only quinacrine DHC in PBS buffer solution, (B) BSA, BSA-bound GSK3 inhibitor and only GSK3 inhibitor in PBS buffer solution. Simultaneous measurements were done at 280 nm and 485 nm wavelengths.

**Table S1:** Dissociation constants, K_d_, from the tryptophan steady-state fluorescence quenching assay.

| Protein | Small molecule | Dissociation constant ($K_d$)<br>$\mu\text{M}$ (micromolar) |
| --- | --- | --- |
| BSA | Fluorescein | 1.7±0.2 |
| BSA | Doxorubicin | 16.5±2.5 |
| BSA | GSK3 inhibitor | 5.2±1.2 |
| BSA | Quinacrine DHC | 4.6±1.1 |
| HEWL | Fluorescein | 3.8±0.6 |
| HEWL | GSK3 inhibitor | 1.7±0.7 |
| HEWL | Quinacrine DHC | 4.7±0.8 |
| Myoglobin | Fluorescein | 3.0±0.5 |
| Myoglobin | Doxorubicin | 18.4±3.1 |
| Myoglobin | GSK3 inhibitor | 7.3±1.8 |
| Myoglobin | Quinacrine DHC | 3.0±0.4 |

**Table S2:** Stokes radius (R, in Å), translational (D_trans_, in µm^2^s^-1^) and rotational diffusion coefficients (D_rot_, in radian^2^/ps) for small molecules and protein crowders computed using HYDROPRO. These values were used as input for BD simulations. D_trans_ and D_rot_ values are for infinite dilution aqueous solution conditions.

| **molecule** | **R** | **$D_{\text{trans}}$** | **$D_{\text{rot}}$** |
| --- | --- | --- | --- |
| doxorubicin | 5.1 | 412 | 9.8E-4 |
| fluorescein | 4.3 | 521 | 2.0E-3 |
| quinacrine | 4.9 | 459 | 1.3E-3 |
| GSK3 inhibitor | 4.5 | 515 | 2.0E-3 |
| HEWL | 15.3 | 109 | 2.1E-5 |
| myoglobin | 16.6 | 102 | 1.7E-5 |
| BSA | 25.7 | 60 | 3.4E-6 |

**Table S3:**
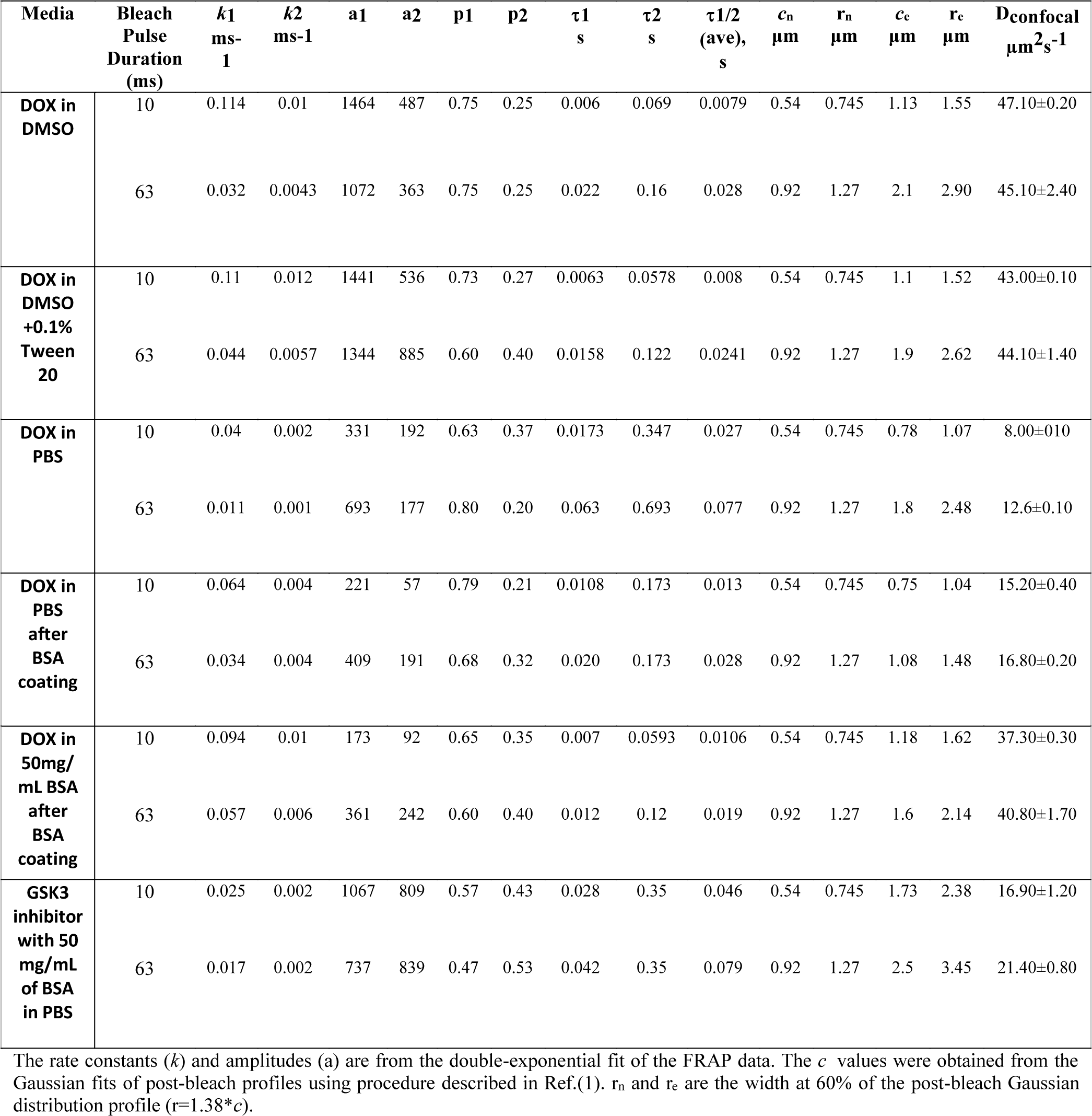
Diffusion coefficients (Dconfocal) of different drugs in PBS buffer or in the presence of protein crowders from measurements with 10ms or 63ms bleach time pulses.

| Media | Bleach Pulse Duration (ms) | $k_1$ ms<sup>-1</sup> | $k_2$ ms<sup>-1</sup> | $a_1$ | $a_2$ | $p_1$ | $p_2$ | $\tau_1$ s | $\tau_2$ s | $\tau_{1/2}$ (ave), s | $c_n$ $\mu\text{m}$ | $r_n$ $\mu\text{m}$ | $c_e$ $\mu\text{m}$ | $r_e$ $\mu\text{m}$ | $D_{\text{confocal}}$ $\mu\text{m}^2\text{s}^{-1}$ |
| --- | --- | --- | --- | --- | --- | --- | --- | --- | --- | --- | --- | --- | --- | --- | --- |
| DOX in DMSO | 10 | 0.114 | 0.01 | 1464 | 487 | 0.75 | 0.25 | 0.006 | 0.069 | 0.0079 | 0.54 | 0.745 | 1.13 | 1.55 | 47.10±0.20 |
|  | 63 | 0.032 | 0.0043 | 1072 | 363 | 0.75 | 0.25 | 0.022 | 0.16 | 0.028 | 0.92 | 1.27 | 2.1 | 2.90 | 45.10±2.40 |
| DOX in DMSO +0.1% Tween 20 | 10 | 0.11 | 0.012 | 1441 | 536 | 0.73 | 0.27 | 0.0063 | 0.0578 | 0.008 | 0.54 | 0.745 | 1.1 | 1.52 | 43.00±0.10 |
|  | 63 | 0.044 | 0.0057 | 1344 | 885 | 0.60 | 0.40 | 0.0158 | 0.122 | 0.0241 | 0.92 | 1.27 | 1.9 | 2.62 | 44.10±1.40 |
| DOX in PBS | 10 | 0.04 | 0.002 | 331 | 192 | 0.63 | 0.37 | 0.0173 | 0.347 | 0.027 | 0.54 | 0.745 | 0.78 | 1.07 | 8.00±0.10 |
|  | 63 | 0.011 | 0.001 | 693 | 177 | 0.80 | 0.20 | 0.063 | 0.693 | 0.077 | 0.92 | 1.27 | 1.8 | 2.48 | 12.6±0.10 |
| DOX in PBS after BSA coating | 10 | 0.064 | 0.004 | 221 | 57 | 0.79 | 0.21 | 0.0108 | 0.173 | 0.013 | 0.54 | 0.745 | 0.75 | 1.04 | 15.20±0.40 |
|  | 63 | 0.034 | 0.004 | 409 | 191 | 0.68 | 0.32 | 0.020 | 0.173 | 0.028 | 0.92 | 1.27 | 1.08 | 1.48 | 16.80±0.20 |
| DOX in 50mg/mL BSA after BSA coating | 10 | 0.094 | 0.01 | 173 | 92 | 0.65 | 0.35 | 0.007 | 0.0593 | 0.0106 | 0.54 | 0.745 | 1.18 | 1.62 | 37.30±0.30 |
|  | 63 | 0.057 | 0.006 | 361 | 242 | 0.60 | 0.40 | 0.012 | 0.12 | 0.019 | 0.92 | 1.27 | 1.6 | 2.14 | 40.80±1.70 |
| GSK3 inhibitor with 50 mg/mL of BSA in PBS | 10 | 0.025 | 0.002 | 1067 | 809 | 0.57 | 0.43 | 0.028 | 0.35 | 0.046 | 0.54 | 0.745 | 1.73 | 2.38 | 16.90±1.20 |
|  | 63 | 0.017 | 0.002 | 737 | 839 | 0.47 | 0.53 | 0.042 | 0.35 | 0.079 | 0.92 | 1.27 | 2.5 | 3.45 | 21.40±0.80 |

The rate constants ( $k$ ) and amplitudes ( $a$ ) are from the double-exponential fit of the FRAP data. The $c$ values were obtained from the Gaussian fits of post-bleach profiles using procedure described in Ref.(1). $r_n$ and $r_e$ are the width at 60% of the post-bleach Gaussian distribution profile ( $r=1.38*c$ ).

**Table S4:** Contacts between small molecules in BD simulations for each of the four compounds in aqueous solution. The average ± standard deviation of the number of small molecule-small molecule contact interactions during three replica BD simulations, each of 10 μs length, is given for each compound. Each simulated periodic system was composed of 80 small molecules in implicit solvent. Interactions were defined as contacting when one or more non-hydrogen atoms of one molecule were within a distance of 4.5 Å of a non-hydrogen atom of another molecule. Interactions were recorded every 500 ps, and the number of contacts was averaged first over time for one simulation, and then over the three replica simulations.

| **Small molecule** | **Number of contact interactions** |
| --- | --- |
| doxorubicin | $0.0312 \pm 0.0002$ |
| fluorescein | $0.014 \pm 0.001$ |
| GSK3 | $0.141 \pm 0.004$ |
| quinacrine | $0.032 \pm 0.001$ |

**Table S5:**
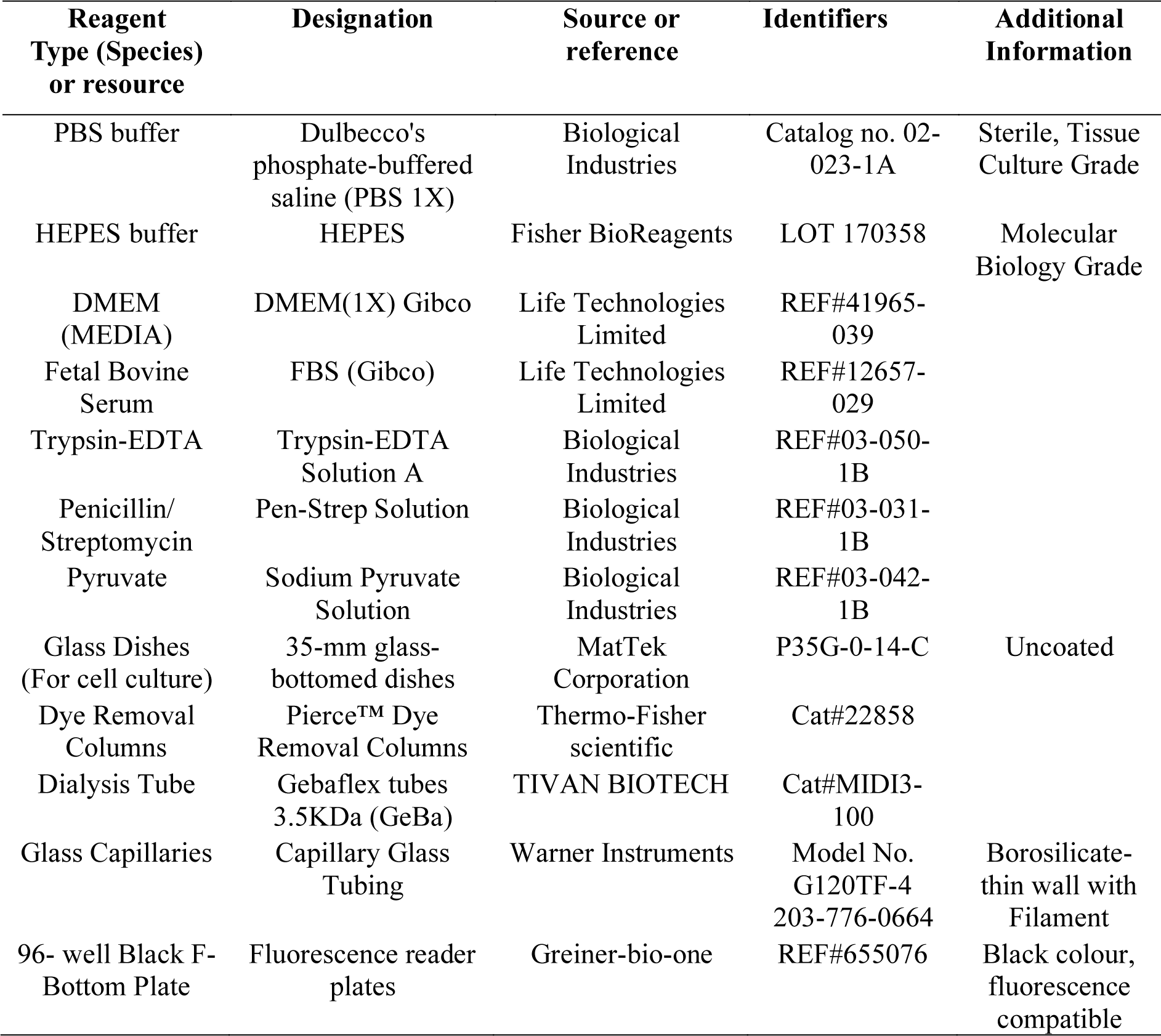
Key resources

**Table S6:** Side lengths of periodic simulation boxes (box, in Å), concentration of protein crowder (in mg/mL), and number (N) of protein crowders and small molecules in each BD simulation.

| **Box length** | **Concentration** | **N protein crowder** | **N small molecule** |
| --- | --- | --- | --- |
| 374 | 0 | - | 80 |
| HEWL |  |  |  |
| 1015 | 10 | 440 | 80 |
| 806 | 20 | 440 | 80 |
| 594 | 50 | 440 | 80 |
| myoglobin |  |  |  |
| 1376 | 5 | 440 | 80 |
| 1092 | 10 | 440 | 80 |
| 867 | 20 | 440 | 80 |
| 638 | 50 | 440 | 80 |
| BSA |  |  |  |
| 1695 | 10 | 440 | 80 |
| 1345 | 20 | 440 | 80 |
| 991 | 50 | 440 | 80 |
| 624 | 200 | 440 | 80 |

**Table S7:** Translational diffusion coefficients (Dtrans, µm^2^s^-1^) from experiments and calculated using HYDROPRO for small molecules in infinite dilution aqueous solution before and after the adjustment of the radius of the atomic elements (AER). Differences are expressed as a percentage of the experimental diffusion coefficient. Adjusting the AER value for small molecules was necessary because the original AER value was parametrized to reproduce experimental results for proteins.The details about HYDROPRO are discussed in the supplemental methods text section (16).

| Small molecule | Experimental $D_{\text{trans}}$ | AER of 2.9 Å<br>(before adjustment) | | AER of 1.2 Å<br>(after adjustment) | |
| --- | --- | --- | --- | --- | --- |
| | | Calculated $D_{\text{trans}}$ | % difference | Calculated $D_{\text{trans}}$ | % difference |
| Benzene | 1090 (2) | 547 | 50 | 987 | 9 |
| Glycine | 1055 (2) | 546 | 48 | 995 | 6 |
| Alanine | 910 (2) | 518 | 43 | 929 | -2 |
| Threonine | 798 (2) | 483 | 39 | 822 | -3 |
| galactose | 666 (3) | 449 | 33 | 726 | -9 |

